# The lncRNA landscape of cardiac resident macrophages and identification of *Schlafenlnc* as a regulator of macrophage migratory function

**DOI:** 10.1101/2022.11.30.518576

**Authors:** Anne Dueck, Lara Althaus, Kathrin Heise, Dena Esfandyari, Seren Baygün, Ralf P. Brandes, Julien Gagneur, Nicolas Jaé, Percy Knolle, Matthias S. Leisegang, Lars Maegdefessel, Thomas Meitinger, Dierk Niessing, Niklas Petzold, Deepak Ramanujam, Hendrik Sager, Christian Schulz, Evangelos Theodorakis, Anna Uzonyi, Tobias Weinberger, Ilka Wittig, Michael Bader, Marc Schmidt-Supprian, Stefan Engelhardt

**Affiliations:** Institute of Pharmacology and Toxicology, Technical University Munich, Munich, Germany; DZHK (German Centre for Cardiovascular Research), partner site Munich Heart Alliance, Munich, Germany; Institute of Experimental Hematology, Central Institute for Translation Cancer Research, School of Medicine, Technical University Munich, Munich, Germany; Institute for Cardiovascular Physiology, Goethe University, Frankfurt, Germany; DZHK (German Centre for Cardiovascular Research), partner site RheinMain, Frankfurt, Germany; Institute of Informatics, Technical University Munich, Munich, Germany; Institute of Human Genetics, Klinikum rechts der Isar, School of Medicine, Technical University of Munich, Munich, Germany; Computational Health Center, Helmholtz Center Munich, Neuherberg, Germany; Institute for Cardiovascular Regeneration, Goethe University Frankfurt, Frankfurt am Main, Germany; Institute for Molecular Immunology, MRI, Technical University Munich, Munich, Germany; Department of Vascular and Endovascular Surgery, Technical University Munich, Munich, Germany; Institute of Structural Biology, Molecular Targets and Therapeutics Center, Helmholtz Munich, Neuherberg, Germany; Institute of Pharmaceutical Biotechnology, Ulm University, Ulm, Germany; Department of Cardiology, German Heart Centre Munich, Technical University Munich, Munich, Germany; Medizinische Klinik und Poliklinik I, Klinikum der Universität München, Ludwig-Maximilians-Universität, Munich, Germany; Max-Delbrück-Center for Molecular Medicine (MDC), Berlin, Germany

**Keywords:** long non-coding RNA, cardiac resident macrophages, pressure overload, *Schlafenlnc*, single cell sequencing

## Abstract

Cardiac resident macrophages (crMPs) were recently shown to exert pivotal functions in cardiac homeostasis and disease, but the underlying molecular mechanisms are largely unclear. Long non-coding RNAs (lncRNAs) are increasingly recognized as important regulatory molecules in a number of cell types, but neither the identity nor the molecular mechanisms of lncRNAs in crMPs are known. Here, we have employed deep RNA-seq and single cell RNA sequencing to resolve the crMP lncRNA landscape from healthy and diseased murine myocardium. CrMPs express previously unknown and highly cell type-specific lncRNAs, among which one lncRNA, termed *Schlafenlnc*, was particularly abundant and enriched in crMPs. We found *Schlafenlnc* to be necessary for migration-associated gene expression in macrophages *in vitro* and *in vivo* and essential for their adhesion and migration. Collectively, our data provide a basis to the systematic characterization of lncRNAs in crMPs and establish *Schlafenlnc* as a critical regulator of macrophage migratory functions.

## INTRODUCTION

Long non-coding RNAs (lncRNAs) are the largest group of non-coding RNA molecules, with 14,764 lncRNA genes currently annotated in mouse and 19,933 in humans (Gencode in October 2022). Although only a small fraction of these has been functionally characterized, there is strong evidence that lncRNAs, through a variety of mechanisms, adopt critical roles in disease, for example in cancer^1^, metabolic, neurologic^2^ and cardiovascular disease^3^. Macrophage lncRNAs in cardiovascular disease have been primarily characterized in the vasculature and within the context of atherosclerosis, e.g., formation of foam cells or polarization (reviewed in ^4^). With regard to cardiac disease, the analysis of lncRNAs and their functions has largely focused on cardiomyocytes^5^, endothelial cells^6^ and fibroblasts^7^ (reviewed in ^3^). By contrast, very little is known about lncRNAs in cardiac resident macrophages (crMPs), although this cell type constitutes the largest population of immune cells in the myocardium during homeostasis and also determines cardiac disease processes (reviewed in ^8^). CrMPs have in recent years been characterized with respect to their developmental origin^9,10^, their composition^10,11^ and their putative functions such as roles in cardiac conduction^12,13^, hypertrophy^14^ or lymph vessel growth^15^. Although it can be readily assumed that many of these functions are subject to lncRNA-mediated regulation, a comprehensive description of the lncRNome from crMPs or functional data on a crMP-specific lncRNA is missing thus far.

In this study, we established the first comprehensive analysis of the lncRNA portfolios of the major cardiac cell types in health and disease, with particular emphasis on macrophages. In mouse myocardium, we identified lncRNAs that are specific for, and define, this cell type. One crMP lncRNA that has previously not been analyzed and which we termed *Schlafenlnc*, stands out for its strong enrichment and high abundance in crMPs. Upon functional characterization of a *Schlafenlnc* knockout in a mouse model for left ventricular pressure overload, we found that this lncRNA regulates macrophage abundance in the heart and the ability of this cell type to exert chemotactic migration. We therefore propose that *Schlafenlnc* critically regulates the function of cardiac macrophages in disease.

## RESULTS

### lncRNAs are important determinants for the definition of specific cell types

To determine the long non-coding RNA (lncRNA) repertoire in the major cardiac cell types and its changes under disease conditions, we subjected mice to chronic left ventricular pressure overload (induced by transverse aortic constriction, TAC), an established model of cardiac inflammation that leads to hypertrophic and fibrotic remodeling of the heart. At day 6 after surgery, we sacrificed the mice, isolated primary cells by Langendorff digestion^16^, and performed droplet-based single cell RNA sequencing (scRNA-seq, Chromium platform) (Fig. 1A). This experimental approach analyzes all cardiac non-myocytes, while the number of cardiac myocytes is decreased due to size constraints of the method. Unsupervised clustering of both samples based on the expression of both coding and non-coding RNAs with the Seurat analysis tool (Seurat V3.2^17^, Gencode vM22^18^) yielded a total of 20,010 cells after quality control. These comprised all major non-myocyte cardiac cell types (Seurat clustering resolution 0.3, Fig. 1B), including subpopulations of immune cells, fibroblasts, endothelial cells and mural cells (Fig. 1B). Differential gene expression analysis from these cell types revealed distinct sets of protein-coding genes that define each major cardiac cell type as individual clusters (Extended Data Fig. S1), in line with previous reports^19,20^.

**Figure 1:**
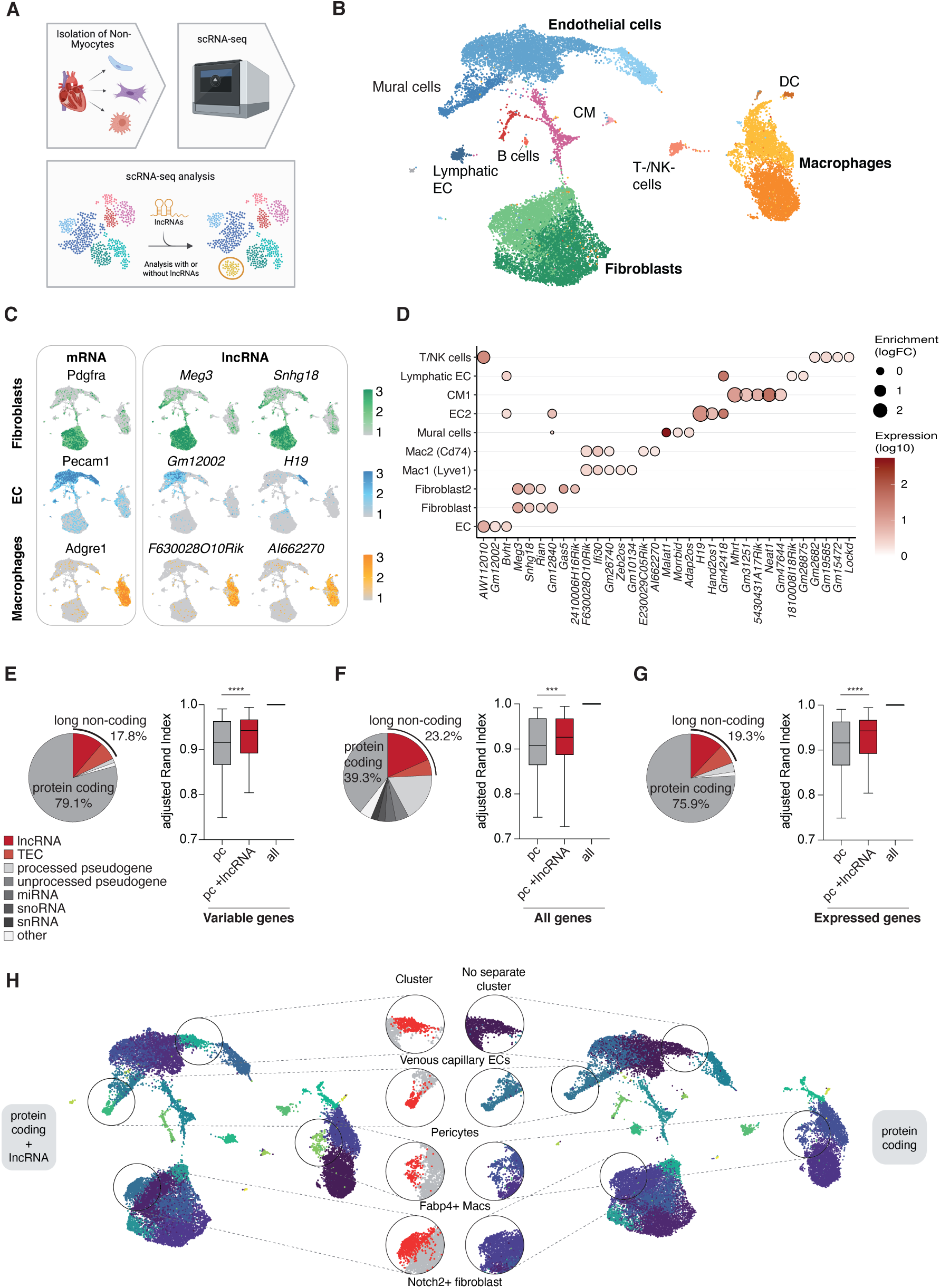
LncRNA are important determinants for definition of specific cell types. (A) Illustration for the measurement of lncRNA in murine myocardium and re-clustering analysis. (B) UMAP projection of murine myocardium as measured by single-cell sequencing (scRNA-seq). Cell populations are colored by type as indicated by the label. DC: dendritic cells, EC: endothelial cells, CM: cardiac myocytes. (C) Bubble plot showing lncRNA marker genes conserved across conditions. Bubble size corresponds to enrichment (logFC) and fill color to expression value (log10). Only values/bubbles with a significant enrichment (p-val < 0.05) are shown. (D) Feature plots showing mRNA and lncRNA markers for fibroblasts, endothelial cells (EC) and macrophages. (E), (F), (G) Adjusted Rand Index (ARI) calculation. In order to compare the clusterings with and without lncRNAs, the respective complete gene set was considered to yield the optimal clustering. We iterated the clustering in 0.01 steps with resolutions ranging from 0.10 to 1.00, testing for the similarity to optimal clustering using adjusted Rand index scoring. Here, the more similar a clustering would be to the “optimal” one the closer it will be to 1. Displayed as boxplot with median and quartiles. (E) ARI calculation for the top variable genes (3,000 genes). (F) ARI calculation for all annotated genes (Gencode vM22^18^). (G) ARI calculation for all expressed genes (expression value >0). pc: protein coding (H) Umap projection of murine myocardium with cell population clustering based on protein coding genes plus lncRNA genes (left panel) or only protein coding genes (right panel). The middle panel shows differentially assigned cell population clusters.

Despite the generally lower expression levels of lncRNAs in comparison to protein-coding genes^21–24^, we were able to readily detect over 440 lncRNAs. Several of the most strongly expressed lncRNAs exhibited a very high level of cell type specificity (Fig. 1C and 1D), suggesting that cardiac cell types can be defined based on their lncRNA transcriptomes. As Fig. 1D shows, our data not only confirmed the previously reported expression of *Meg3* in fibroblasts^25^, *H19* in endothelial cells^26^ and *Mhrt* in cardiac myocytes^27^, but also revealed lncRNAs that had hitherto not been reported to be cell type-specific in the heart. These lncRNAs may reasonably be presumed to potentially exert important roles in their specific cell type, prompting for functional characterization.

To validate the contribution of lncRNAs to cell type identity, we performed differential clustering of the scRNA-seq data sets with or without lncRNA genes, and tested for clustering precision. Independent of whether the top 3,000 variable genes defined by the Seurat pipeline, all annotated genes (n=55,487) or expressed genes (n=19,719, expression value >0) were used to perform clustering, the inclusion of lncRNA marker genes significantly improved clustering compared to protein coding genes alone (Fig. 1E-G). Compared to clustering based on protein-coding genes alone, this resulted in an overall increased sensitivity (indicated by an increased adjusted Rand Index value) and yielded additional subclusters of cell types. Specifically, the detection of clusters representing venous capillary endothelial cells, pericytes, *Fabp4*^+^ macrophages and *Notch2*^+^ fibroblasts was dependent on the inclusion of lncRNAs (resolution 0.65) (Fig. 1H, Extended Data Fig. S1B).

In conclusion, lncRNAs were shown to be a valuable determinant of defining clusters and fine-tuning cell type assignations.

As a means to approach lncRNAs of potential disease relevance, we next thought to assess the expression of lncRNAs during cardiac disease by comparing the datasets obtained from sham-treated vs. TAC-treated mice. Upon increasing the clustering resolution from 0.3 to 0.7, a more detailed and granular cell assignment of lncRNA genes to cellular subtypes was achieved (Extended Data Fig. S2A and S2B). With this setup, we performed differential gene expression analysis for the sub-populations of the major cell types in the heart, and visualized the top de-regulated lncRNAs (Fig. 2A-C, Extended Data Fig. S2C). Some lncRNAs were regulated after TAC throughout all subpopulations of a cell type, e.g., Gm29686 in fibroblasts (Fig. 2A), whereas others were only significantly regulated within a defined subpopulation, e.g., Gm49785 in the fibroblast subpopulation (FB6) (Fig. 2A). This subpopulation of fibroblasts represents activated myofibroblasts (high expression of periostin), a cell type involved in extracellular matrix secretion and immune cell recruitment^28^. Similarly, several lncRNAs displayed dysregulation within cardiac endothelial cells (Fig. 2B) and cardiac macrophages (Fig. 2C), resolved to the level of cellular subtypes.

**Figure 2:**
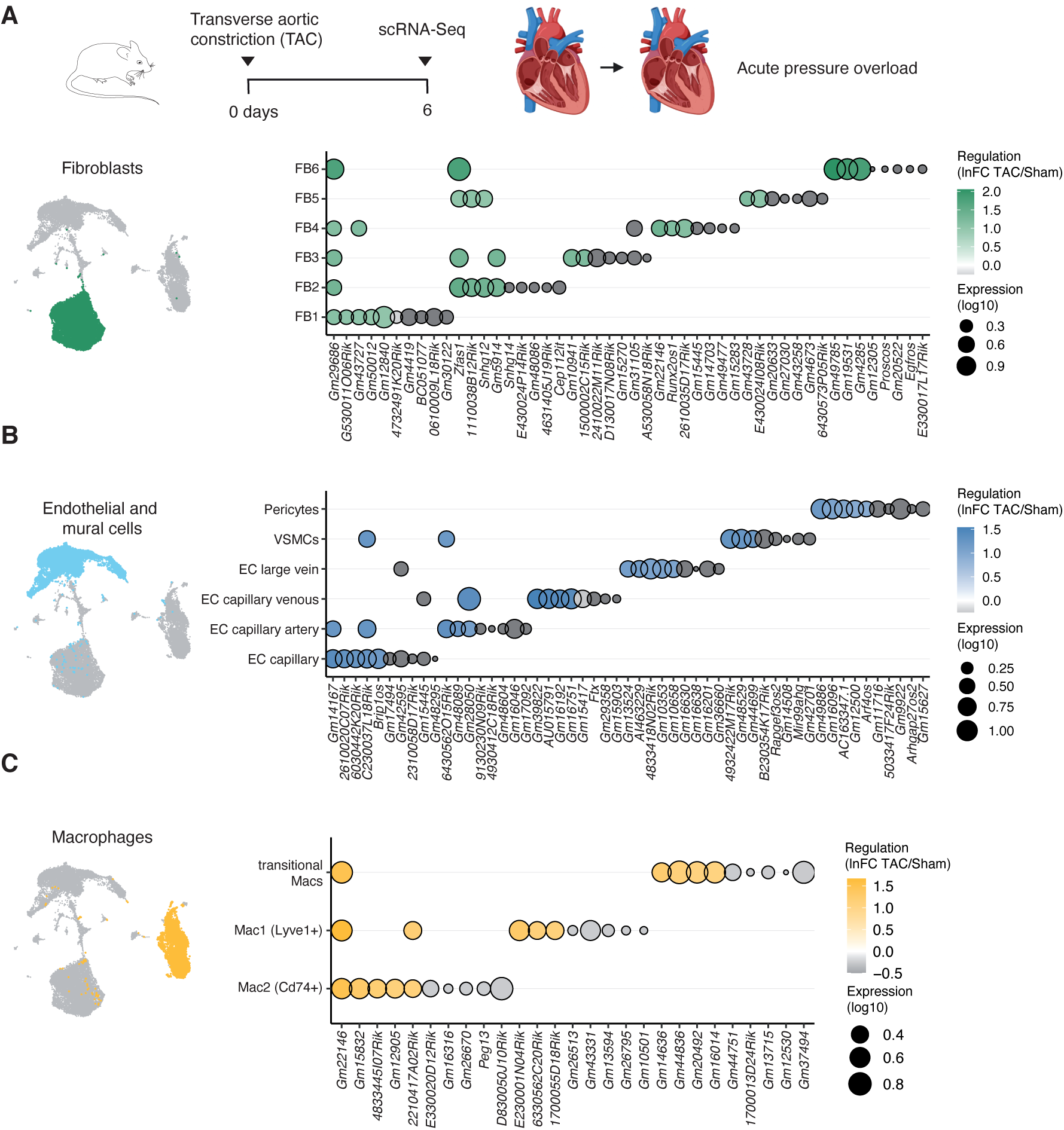
Dynamic changes of lncRNA expression in crMPs during pressure overload. Bubble plot showing the top 5 up-regulated and top 5 down-regulated lncRNA genes after TAC for the sub-populations of the major cell types in the heart: (A) Fibroblasts (B) Endothelial cells and (C) Macrophages. Fill color corresponds to regulation in TAC vs. sham (lnFC TAC/sham) and bubble size corresponds to expression value (log10). Only values with a significant enrichment (p-val < 0.05) are shown. Some genes are among the top regulated genes in more than one cell type.

Of particular interest are lncRNAs in macrophages of the myocardium, since no lncRNA had thus far been identified or functionally characterized in this cardiac cell type. As a first approach to the potential function of the lncRNAs we had identified as deregulated in crMPs in the TAC model, we correlated these lncRNAs with biological processes of their overlapping neighboring genes (Extended Data Fig. S2D). Since lncRNAs are often co-expressed and co-regulated with proximal genes in *cis*^29^, we speculated that the latter may give insight into the biological context within which these lncRNAs function. For example, genes antisense to downregulated lncRNAs in the Mac1 cluster (*Lyve1* macrophages) after TAC are associated with programmed cell death processes and genes antisense to downregulated lncRNAs in the Mac2 cluster (*Cd74*+ macrophages) after TAC were predominantly linked to adhesion pathways (Extended Data Fig. S2D), suggesting that the reported de-regulated lncRNAs might also be involved in the regulation of these processes.

Taken together, we report a comprehensive inventory of regulated lncRNA genes in the major non-myocyte cell types of the heart during pressure overload. These lncRNA genes will be interesting candidates to be studied in more detail.

### Determination and annotation of the lncRNA landscape of cardiac resident macrophages

As the sequencing depth of current single cell sequencing methodology limits detection sensitivity, we therefore complemented our scRNA-seq data by performing deep RNA-Seq on basal total murine myocardium as well as crMPs that were isolated using well-established cell markers (CD45, CD64, CD11b and F4/80^9,10^). These surface proteins collectively labelled essentially all crMPs in our scRNA-seq data (Extended Data Fig. S3A), corroborating our purification strategy. Using a pipeline to identify and quantify novel lncRNAs (Extended Data Fig. S3B), we identified 250 genes that had previously not been annotated by Gencode vM22^18^. The identified novel lncRNAs showed the same characteristics as the known lncRNAs with respect to having no coding probability (true non-coding) and transcript length (Extended Data Fig. S3C upper panel), but were more highly expressed in our data set than the known lncRNA population (Extended Data Fig. S3C lower panel). RNA-seq from fractionation experiments showed for known lncRNAs a distribution of 80% in the nucleus (and 20% to the cytoplasm), the novel lncRNAs are slightly less nuclear in localization (70% nuclear, 30% cytoplasmic; Extended Data Fig. S3D).

As a further approach to the function of the newly identified lncRNAs, we built on the observation that the products of lncRNA genes and protein-coding genes which are in proximity of 5kb or less and in antisense overlapping orientation are prone to interact in the same biological processes^30^. The distribution of the interaction categories (e.g. “antisense overlapping”) was comparable between novel and known lncRNAs (Extended Data Fig. S3E). Notably, 15 novel lncRNA genes were found without any interaction, categorizing them as novel long intergenic RNAs (lincRNA). For both known and novel lncRNAs we found many well-expressed genes (> 5 transcripts per million reads, TPM) that are specific in cardiac macrophages compared to total murine myocardium (containing myocytes, endothelial cells, fibroblasts, and macrophages) (Fig. 3A).

**Figure 3:**
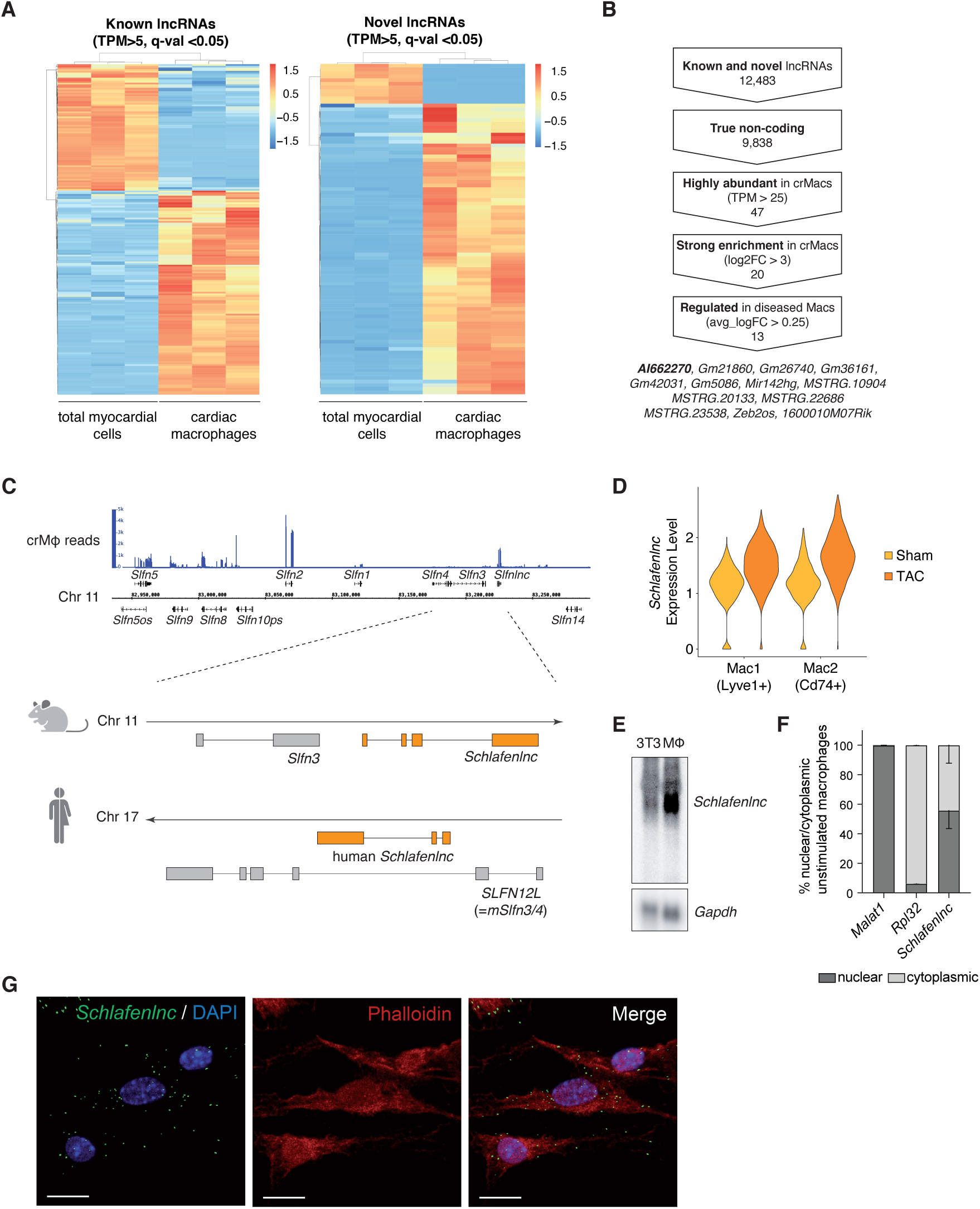
Determination and annotation of the lncRNA landscape of cardiac resident macrophages. (A) Unsupervised clustering of known and novel lncRNAs using normalized counts values, rows were averaged across samples for clustering. Genes shown are expressed with a TPM value >5 and differentially regulated (enriched) with a q-val of <0.05. (B) Flow scheme for the identification of lncRNA candidates using the indicated filters. (C) The Schlafen family locus showing the expression in cardiac resident macrophages (accumulated plus and minus strand). Lower panel: Zoom in of the *Schlafenlnc* locus in mouse as well as human for comparison. (D) Expression of the lncRNA *Schlafenlnc* in Mac1 (*Lyve1*+ macrophages) and Mac2 (*MHCII* high, *Cd74*+ macrophages) after sham or TAC in the scRNA-seq data. (E) Northern Blot showing the expression of *Schlafenlnc* transcripts in bone marrow-derived macrophages in comparison to fibroblasts (NIH3T3 cell line). (F) Relative nuclear and cytoplasmic localization of *Malat1* (nuclear lncRNA), *Rpl32* (mRNA) and *Schlafenlnc* as determined by fractionation of macrophages followed by quantification through RNA-Seq. Data are normalized counts displayed as mean with SEM. (G) Detection of *Schlafenlnc* (green) in macrophages by RNA *in situ* hybridization (RNAScope). Nuclei were stained with DAPI (blue), Phalloidin (red) was used for outlining the cellular body (Scale bars = 15µm).

### Identification of lncRNA *Schlafenlnc* as an abundant and enriched macrophage lncRNA

In order to select lncRNA candidates for experimental validation, we further filtered the datasets from scRNA-Seq data and RNA-Seq based on the following stringent criteria: low coding probability according to CPAT score (true non-coding RNA < 44% coding probability), absolute expression > 25 TPM in macrophages, enrichment of log2FC >3 and FDR <0.05, and de-regulation in TAC (scRNA-seq data) FC >0.25). This yielded 13 lncRNA candidates that are highly abundant, enriched and regulated in cardiac macrophages (Fig. 3B).

We found one of these candidates, lncRNA *AI662270*, particularly interesting due to its genomic association with a cluster of eight genes on chromosome 11 which encode members of the Schlafen family of proteins (Fig. 3C). Schlafen proteins are found in human and mouse and known to regulate important processes and functions such as immune cell development, invasion and migration^31^. *AI662270*, however, is a hitherto uncharacterized gene within this locus. The genomic location of lncRNA *AI662270* in the Schlafen cluster led us to name the transcript *Schlafenlnc*.

The Schlafen locus of protein-coding genes is very similar between human and mouse in general, yet the internal order of the proteins (and lncRNAs) is slightly different. A study by Ulitsky and colleagues developed a pipeline to identify syntenic genes across multiple organisms, where lncRNAs are tested on the relative location and orientation to orthologous protein-coding genes^32^. By comparing Gencode annotations^18^ of human (V30) and mouse (vM21) with their pipeline, the gene *ENSG00000267074* was designated as the putative human *Schlafenlnc* homolog (Fig. 3C). It is expressed in various human cell types of leukocyte origin (using the RNA-Get module Encode data sets^33^, Extended Data Fig. S4A), making it an attractive molecule for studying the observed phenotypes in human cells. Besides the analysis of expression, we tested if the two lncRNAs share common potential binding partners using a pretrained deep neural network (DNN)^34^ on eCLIP-seq data^35^. Indeed, the mouse *Schlafenlnc* and human *ENSG00000267074* contain three major binding modules of the same or similar proteins (Extended Data Fig. S4B and S4C), supporting the hypothesis of conservation.

Murine *Schlafenlnc* was up-regulated after TAC in both macrophage populations, but even more pronounced in Mac2 (*MHCII*-high, *Cd74*+) macrophages (Fig. 3D). Its strong expression made it readily detectable by Northern blotting (Fig. 3E). Interrogation of the RNA-seq datasets from the nucleus and the cytosol, as well as RNA *in situ* hybridization, showed that *Schlafenlnc* localizes to both compartments (Fig. 3F and G, Extended Data Fig. S4D).

### *Schlafenlnc* determines myocardial macrophage infiltration and polarization in response to pressure overload

To assess the phenotype of a macrophage-specific deficiency of *Schlafenlnc in vivo*, we generated a *Schlafenlnc-*deficient mouse line (*Schlafenlnc^-/-^*) by CRISPR/Cas9 and examined these mice under basal and disease conditions. The dominant transcript was targeted with specific sgRNAs, including the promoter region upstream of *Schlafenlnc* (determined by H3K4me3 signals in murine bone marrow-derived macrophages, ENCODE data set ENCSR000CFF^36^) to abrogate all transcription from that locus (Extended Data Fig. 5 A). Deletion was verified by PCR from genomic DNA (Extended Data Fig. 5 B). The early compensation stage of the TAC model is characterized by sterile inflammation and both an influx of monocytes as well as proliferation of resident macrophages. We therefore performed sham and TAC surgery (Fig. 4 A) and analyzed the heart phenotype and monocyte/macrophage numbers after 6 days (Fig. 4 B). The heart weight of *Schlafenlnc*-deficient mice was significantly reduced (Fig. 4 C) and heart function (comparing ejection fraction of sham and TAC mice) was significantly improved (Fig. 4 D). Furthermore, *Schlafenlnc^-/-^* hearts showed a substantial decrease in pressure overload-induced interstitial fibrosis (Fig. 4 E) and interestingly a significantly reduced monocyte/macrophage infiltration after TAC, as determined by immunostaining of left ventricular myocardial sections against the monocyte/macrophage marker CD68 (Fig. 4 F).

**Figure 4:**
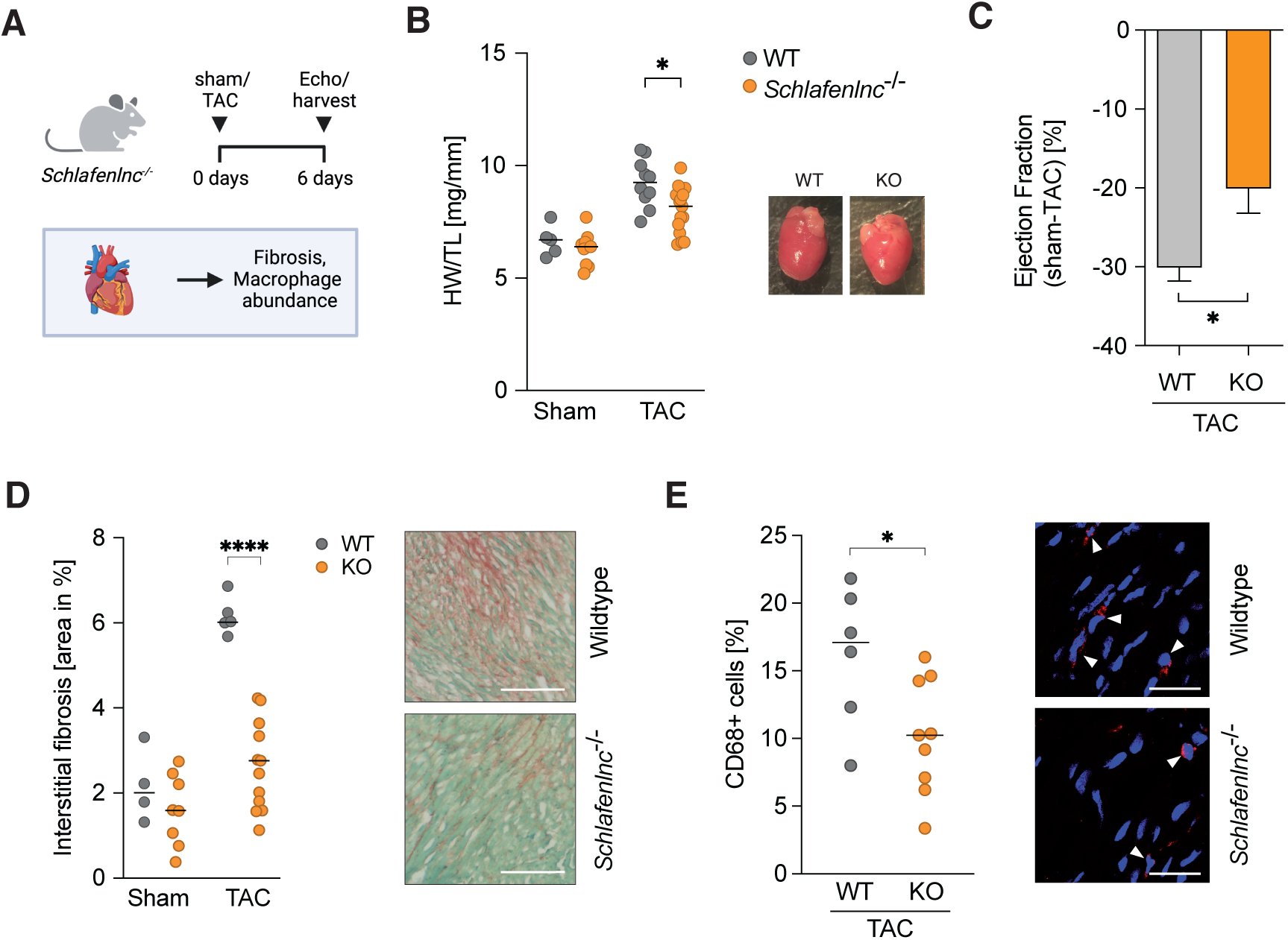
***Schlafenlnc*-deficient mice show improved cardiac function and reduced inflammatory response after transverse aortic constriction (TAC).** (A) Experimental setup of cardiac phenotyping of *Schlafenlnc^-/-^* and wildtype mice. (B) Quantification of normalized heart weight after TAC. HW= heart weight, TL= tibia length. Black line = median. C) Ejection fraction changes after 6 days TAC. Data displayed as mean with SEM. (C) Left: Quantification of interstitial fibrosis by Sirius red/Fast green staining. Black line = median. Right: Representative images of interstitial fibrosis, scale bar = 200µm. (D) Representative images and quantification of macrophage/monocyte content by CD68 staining in WT and *Schlafenlnc^-/-^* hearts after 6d after TAC. Black line = median. White arrows label CD68-positive cells, scale bar = 25µm.

To assess the composition of immune cells in more detail we next performed scRNA-seq of the CD45+ cell fraction isolated from the myocardium and blood (Fig. 5 A). As expected, macrophages are the dominant cell population within the heart whereas the blood contains more monocytes (Fig. 5 B). Activated macrophages and monocytes release signals in an auto- and paracrine fashion^19^. *Schlafenlnc^-/-^*mice showed decreased signaling between almost all macrophage and monocyte populations after 6 days of TAC (Fig. 5 C), with a significant decrease in inflammatory signaling pathways such as Osteopontin (*Spp1*) and Macrophage Inhibitory Factor (*Mif*) signaling (Fig. 5 D), both are important factors in recruiting monocytes and other inflammatory cells from the blood after cardiac injury^37,38^. Performing RNA velocity analysis of cardiac leukocytes after TAC (Fig. 5 E) showed gene dynamics of *Schlafenlnc* WT *Ly6C*-high monocytes towards non-classical monocytes as well as a moderate dynamic from *MHCII*-high macrophages to *Lyve1* macrophages. In *Schlafenlnc^-/-^*, the dynamics between different monocyte states was largely abolished while transition dynamics from monocyte-derived macrophages (*MHCII*-high) to resident-like macrophages (*Lyve1*) was strongly increased (Fig. 5 E). Taken together, the analysis of *Schlafenlnc^-/-^* myocardium indicated stronger polarization of cardiac macrophages towards the *Lyve1* macrophage population, which is generally believed to play a more anti-inflammatory role in disease.

**Figure 5:**
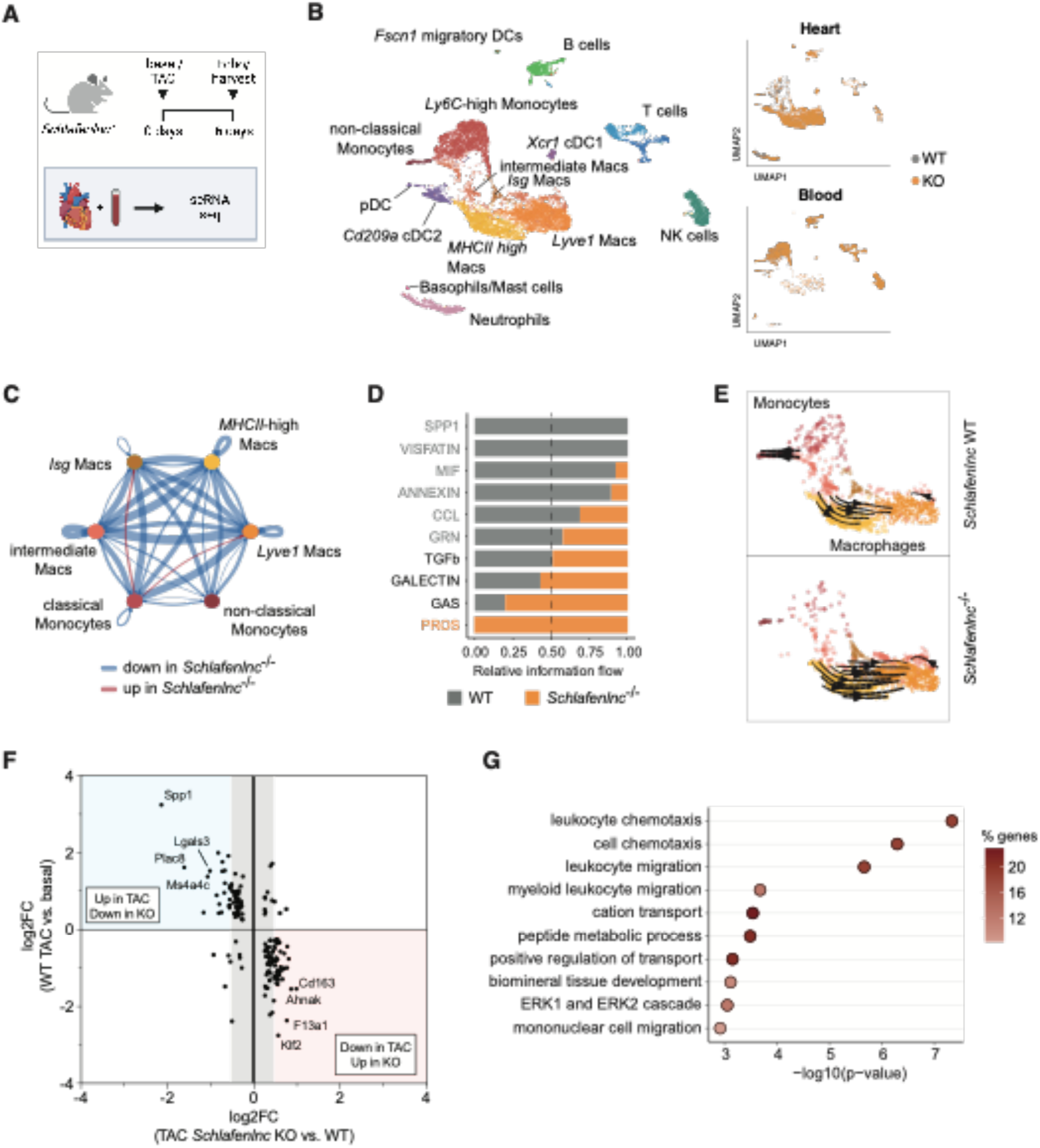
scRNA-seq of *Schlafenlnc^-/-^*mice reveals macrophage polarization towards an anti-inflammatory phenotype after TAC. (A) Experimental setup of scRNA-seq of *Schlafenlnc^-/-^* and wildtype mice. (B) Dimensional reduction (Umap) representation of combined measured cells with assigned cell types. (C) Cell interaction analysis of *Schlafenlnc^-/-^*and wildtype mice (*Cd45*+ cells from blood and heart) after TAC using CellChat^74^. (D) Differential signaling in *Schlafenlnc^-/-^*and WT macrophage and monocyte populations after TAC. Light grey and orange highlighted pathways are significantly enriched in wildtype or *Schlafenlnc^-/-^*, respectively (p-val<0.05). (E) RNA Velocity of *Schlafenlnc^-/-^* and WT cells from heart after 6 days of TAC. Shown are macrophage and monocyte cell clusters. (F) Comparison of de-regulated genes (q-val<0.05) in TAC vs. basal condition in WT compared to de-regulated genes after 6d TAC KO vs. WT in *Lyve1^+^* macrophages (G) Gene ontology terms of genes de-regulated in *Schlafenlnc*-deficient mice and after 6 days TAC.

Next, we asked which genes were changing globally in *Lyve1* macrophages after TAC and upon *Schlafenlnc* deletion. Among the genes that were de-regulated during TAC (WT TAC vs. sham), 64 genes were additionally strongly de-regulated in *Schlafenlnc^-/-^* vs. *Schlafenlnc* WT *Lyve1* macrophages (|Log2FC|>0.5, FDR<0.05). 17 of these genes are classified as coding for secreted proteins (Fig. 5 F, categorization according to Human Protein Atlas^39^), for example, *Lgals3* (Galectin-3) and *Spp1* (Osteopontin), which were de-regulated in all macrophage populations (Extended Data Fig. 5 C). An analysis of the associated biological processes of de-regulated genes in *Lyve1* macrophages (Fig.5 F) showed leukocyte chemotaxis and migration as the strongest impacted pathways (Fig. 5 G).

Taken together, *Schlafenlnc* deletion in immune cells of the heart altered the cardiac macrophage response to early pressure overload towards an anti-inflammatory phenotype and disease-associated phenotypes such as fibrosis and heart weight were improved.

### *Schlafenlnc* is required for macrophage adhesion and migration

To elucidate the function of *Schlafenlnc* in more detail and explore its potential role in migration processes, we generated *Schlafenlnc*-deficient macrophages from Hoxb8-FL macrophage progenitors^40,41^ (Extended Data Fig. S6A). Gene deletion was validated by RNA-Seq (Extended Data Fig. S6A, middle track). As a genomic deletion might have an impact on nearby genes due to a mechanism in *cis* by the deleted lncRNA or due to the deletion of a regulatory element within the DNA, the surrounding region of *Schlafenlnc* (0.5 Mbp in each direction) was analyzed for de-regulation (Extended Data Fig. S6B). 9 of 64 genes were found to be moderately dysregulated (with |log2FC| > 1, FDR < 0.05) in the vicinity of *Schlafenlnc* revealing that there was not a general activation or de-activation of that locus by *Schlafenlnc*. To identify major perturbations on gene expression level, *Schlafenlnc*-deficient monoclonal cell lines (2 clones, each measured in n=4) were subjected to RNA-Seq. These samples were compared to control monoclonal cell lines (two clones with n=3 each) that were generated using an unrelated sgRNA (sequenced as wildtype at the *Schlafenlnc* locus). We observed 2,660 genes significantly de-regulated (|log2FC| > 1, FDR < 0.05) (Fig. 6 A), among those *Schlafenlnc* that dropped from 67 TPM to 0.33 TPM. Among the downregulated genes, we observed an enrichment of genes involved in migration and adhesion similarly to our *in vivo* study (Fig. 6 B, e.g., *Mmp14* and *Lgals3*). Lower amounts of *Lgals3* could not only be observed in our RNA-Seq data but were validated on protein level (Galectin-3 is the gene product of *Lgals3*) by measurement of cell culture supernatant using an enzyme-linked immunosorbent assay (ELISA) (Fig. 6 C). More generally, gene ontology analysis of all significantly de-regulated genes implicated pathways such as regulation of cytokine production, regulation of cell adhesion and chemotaxis that are promoted by *Schlafenlnc* (Fig. 6 D). To validate these results, we performed an adhesion assay by rigorous washing (Fig. 6 E) as well as tested the chemotactic migration ability of *Schlafenlnc*-deficient macrophages with a gradient of the chemokine C5a (Fig. 6 F). Adhesion was diminished by approximately 23% (Fig. 6 E) and chemotaxis was about 50% reduced (Fig. 6 F) in *Schlafenlnc* KO cells compared to wild type controls.

**Figure 6:**
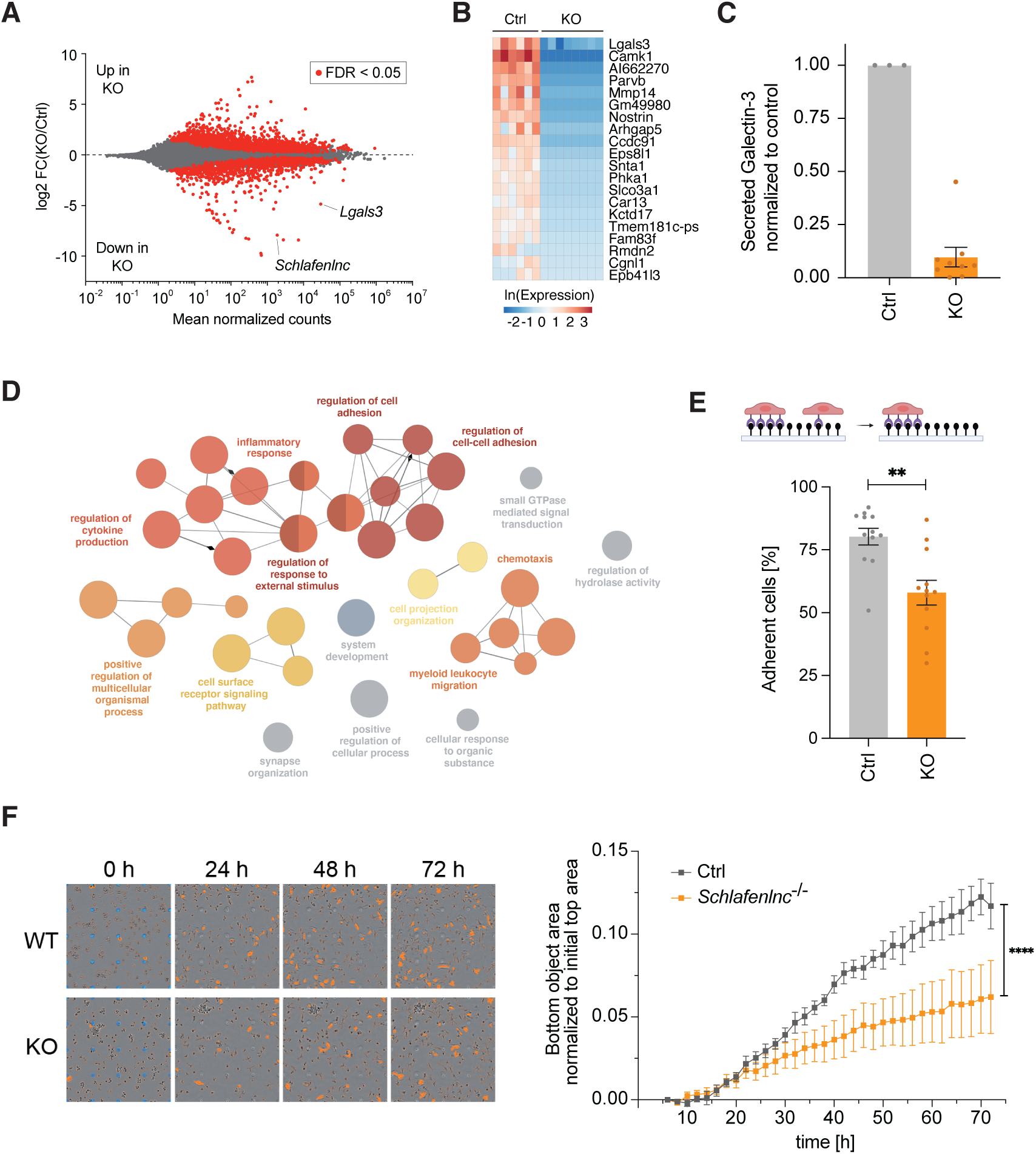
***Schlafenlnc* is required for the activation of macrophage gene programs for cell adhesion and migration.** (A) MA plot showing differentially expressed genes in *Schlafenlnc^-/-^* vs. Ctrl macrophages. Significantly deregulated genes (FDR < 0.05) are shown in red. (B) Heatmap of an unsupervised clustering showing the top 20 downregulated genes in *Schlafenlnc^-/-^* macrophages. (C) Measurement of Galectin-3 protein secretion by control and *Schlafenlnc^-/-^*macrophages as determined by ELISA. The supernatant of one control cell line and three *Schlafenlnc^-/-^* clones were used in n=3 independent experiments. Data points are displayed as mean ±SEM. (D) Functionally grouped gene ontology term analysis of biological processes associated with significantly downregulated genes (FDR < 0.05, log2FC < −1) in *Schlafenlnc^-/-^*macrophages. (E) Adhesion assay with control and *Schlafenlnc^-/-^*macrophages (Ctrl clones n=2, KO clones n=3; 4 independent experiments). Data points are displayed as mean ±SEM. (F) Chemotaxis assay against a gradient of the chemokine C5a with control and *Schlafenlnc^-/-^* macrophages stimulated with 10ng/ml LPS overnight (Ctrl clones n=1, KO clones n=3; 3 independent experiments). Data points are displayed as mean ±SEM.

To identify interaction partners of Schlafenlnc that might contribute to this observed phenotype, we performed a streptavidin pulldown using lysates from bone marrow-derived macrophages incubated with a biotin-labelled Schlafenlnc RNA probe (Extended Data Fig. S6C). The subsequent mass spectrometry analysis identified 27 proteins which significantly bound to Schlafenlnc, with the top enriched proteins being PURA and PURB (Extended Data Fig. S6D). PURA (purine-rich element binding protein A) and PURB (protein B) are Pur family members essential for DNA replication, gene expression, and cell growth (reviewed in ^42^). Both bind purine-rich DNA and RNA sequences, thereby modulating transcription and mRNA stability, can function independently or as a heterodimer^43,44^ and are also known to interact with lncRNAs to regulate gene expression^45^. Next, we validated the interaction with PURA using a specific monoclonal antibody against PURA^44^ and indeed could show significant enrichment of Schlafenlnc after immunoprecipitation of PURA (Extended Data Fig. S6E and F). As a known transcriptional regulator, PURA might therefore be involved in the gene expression changes caused by Schlafenlnc, but further studies are required to elucidate this mechanism.

Together these data obtained on *Schlafenlnc*-deficient macrophages corroborate a cell autonomous role for this lncRNA in macrophages, in line with its specific expression in this cell type. *Schlafenlnc* is required for the activation of macrophage gene programs underlying cell adherence and migration providing a mechanistic basis for the phenotype observed *in vivo* in *Schlafenlnc*-deficient mice.

## DISCUSSION

This study provides the first comprehensive lncRNome of cardiac resident macrophages. One of the most highly enriched and abundant lncRNAs in cardiac resident macrophages – lncRNA *AI662270*, which we termed *Schlafenlnc* – was shown to be involved in adhesion and migration. As for *Schlafenlnc*, a particular characteristic of lncRNAs in general is their high degree of cell type-specificity^21–23^ which makes them important for processes such as development, differentiation of cell types, cellular fate and function (reviewed in ^3^). This characteristic could therefore make lncRNAs the natural choice when describing cell population expression signatures, especially in single cell RNA sequencing (scRNA-seq). scRNA-seq allows for an unbiased approach to identify main- and sub-populations of cells, yet little focus has been set to expressly include lncRNA marker genes. Initially, methodical problems such as library preparation, sequencing depth or analysis software hampered the detection of lncRNAs in such data sets. LncRNAs are still not as readily measured as mRNAs due to their on average lower expression levels, but together with improved scRNA-seq depth and improved annotations, our data suggest that lncRNAs are underestimated markers of cell type definition. Emerging platforms measuring RNA transcripts in tissue such as spatial transcriptomics or *in situ* sequencing (reviewed in ^46^) might be able to leverage that potential further.

Focusing on lncRNAs and their potential regulation in ventricular myocardium during disease, we could identify an extensive set of regulated lncRNAs for all major cell populations. In general, we find more up-regulated than down-regulated genes upon induction of disease, which we believe to be due to the detection limit of scRNA-seq concerning very lowly expressed genes^47^. LncRNAs more specific to and regulated in a subset of cell types might offer new possibilities for therapeutic intervention and to gain insight into regulatory mechanisms specific for defined cell types. As we have shown for the three macrophage clusters (Mac1 (*Lyve1*+), Mac2 (*Cd74*+) and transitional macrophages), the different sets of regulated lncRNAs are implicated in distinct gene ontology pathways, which points to distinct functions of these macrophage populations.

Cardiac resident macrophages have been shown to be key players in diseases such as heart failure and myocardial infarction (reviewed in ^48–51^). In the mouse model of left ventricular pressure overload, crMPs are proliferating during the compensation phase (from 0 days to about 7-14 days). An infiltration of monocyte-derived MPs starts early in disease and increases in the late, decompensated phase^52^. The infiltration of monocyte-derived MPs is typically associated with a long-term detrimental cardiac function^52–59^. This recruitment of monocytes is at least to some degree dependent on crMPs^60^, and suppression of this communication axis might be beneficial. Galectin-3, for example, is known to be increased in heart failure patients^61^ and is associated with murine cardiac remodeling during pressure-overload^62^. It is well known to be involved in the monocyte as well as macrophage migration and infiltration in cardiovascular diseases^63–65^. Our combination of unbiased scRNA-seq measurements and in-depth coverage by RNA-Seq allowed us to identify the hitherto uncharacterized lncRNA *Schlafenlnc* as a regulator of macrophage adhesion and migration. *Schlafenlnc* is up-regulated after 6 days of TAC as well as other genes involved in adhesion and migration such as Galectin-3 and its deletion impacts Galectin-3 expression both *in vivo* and *in vitro*. Our quantitative assessment of macrophage/monocyte numbers (CD68 staining) and the analysis in relative immune cell composition by scRNA-seq after 6 days of TAC point to a mechanism of defective signaling by *Schlafenlnc*-deficient crMPs to initiate recruitment of inflammatory cells such as monocytes. Given its selective expression in crMPs, we believe that targeting of *Schlafenlnc* warrants further exploration as a potential anti-inflammatory therapeutic strategy.

Similar to its orthologue in mice, human *Schlafenlnc* is specific for immune cell types (Extended Data Fig. S4A). In support of conserved function in immune cells, our protein interactor analysis of both murine and human *Schlafenlnc* using deep learning showed similar binding modules for RNA binding proteins (Extended Data Fig. S4B and S4C). Further studies will be required to dissect molecular interactors for *Schlafenlnc* in mouse and human.

Taken together, our study demonstrates that lncRNAs act as critical contributors to the transcriptome-based analysis of cell clusters in line with their crucial role in development and cell fate determination^3^, leading to the definition of cell identities. The combination of single cell sequencing and RNA-Seq approaches led us to the identification of multiple novel candidate lncRNAs that warrant further analysis in cardiac homeostasis and disease. One of the most abundant and macrophage-enriched lncRNAs, *Schlafenlnc*, acts as a crucial regulator of macrophage adhesion and migration.

## METHODS

### Experimental animals, surgery conditions

Surgeries on mice were performed according to the guidelines and regulations of the responsible authorities. Consent was given by the institutional review board of the Regierung of Oberbayern (ROB-55.2-2532.Vet_02-19-194).

Wild type male mice of C57BL/6N background were purchased from Jackson Laboratories and submitted to transverse aortic constriction (TAC) or sham procedure as reported previously^16^ at an age of 8-10 weeks.

To produce *Schlafenlnc* -knockout mice, two sgRNAs (5’ cut site: 5’-GAGGTTCCATCAGTCTGACA-3’, 3’ cut site: 5’-ACATGTCTTTGGTAGGAGCA-3’) targeting the flanking region of the *Schlafenlnc* gene in the mouse genome were designed and synthesized by IDT (Coralville, IA, USA). A mixture of Cas9 protein (IDT) and sgRNAs was electroporated into C57BL/6 mouse zygotes. Genomic DNA of newborn mice was amplified using the primers Slfnlnc1 (5’-TTCCTTCAGAGGTCCATCAG-3’) and Slfnlnc4 (5’-AGCCAGTCTAAAGACTCTGG-3’). The appearance of a 360 bp band identified founder mice with a complete loss of the *Schlafenlnc* gene. The animals were bred to homozygosity and used for the experiments.

Echocardiography was performed as reported before^16^.

### Primary cell isolation from adult murine heart

Hearts of C57BL/6N mice were harvested at an age of 8-10 weeks and subjected to digestion by perfusion via the coronary arteries using Collagenase II (Worthington, Lakewood, NJ) as described previously^66^. Cell suspension was processed according to subsequent protocol/experiment.

### Single cell cloning and sequencing

Primary cells from a sham^16^ and an adult mouse heart 6d after TAC were washed three times with PBS to remove cell debris and passed successively through 100µm, 70µm and 30µm filters. The single cell library was generated using the Chromium Single Cell 3ʹ GEM, Library & Gel Bead Kit v2 (10X Genomics) with a target of 10,000 cells per sample. The single cell library was cloned according to the manufacturer’s instructions. Sequencing was performed on a HiSeq4000 (Illumina).

Primary cells from the hearts from *Schlafenlnc^-/-^*and littermate wildtype mice in basal condition as well as after 6d TAC were passed through a 100µm filter and centrifuged for 1min at 100*xg* to pellet cardiomyocytes. The supernatant (non-myocytes) was passed through a 40µm filter and washed twice in magnetic-activated cell sorting (MACS) buffer (PBS + 0.5% BSA + 2mM EDTA) to result in 200µl cell suspension. For the isolation of PBMCs, blood was diluted with 5 ml of MACS buffer and layered onto 5ml of Ficoll-Paque (17144002, Cytiva), followed by centrifugation at 400xg and 4°C for 30min in a swinging rotor without a brake. The white layer containing the cells was aspirated, transferred to a 50ml tube and mixed with 30ml of MACS buffer. After centrifugation for 5 min at 300xg and 4°C, the pellet was washed once more with 30ml MACS buffer (200xg and 4°C for 5min) and resuspended in 200µl

MACS buffer. Cardiac and blood leukocytes (basal condition) were further purified using MACS, labelled with cell hashing antibodies and isolated by FACS (all steps at 4°C): Cell suspensions were incubated with Fc-block (1:50, BD Pharmingen 553142) for 10min and subsequently incubated with mouse CD45 MicroBeads (Miltenyi Biotec) for 15min. Cells were washed with MACS buffer and MACS was performed using an AutoMACS (Miltenyi Biotec) running the program *possel_s*. After elution, pelleted cells were resuspended in MACS buffer containing anti-CD45-FITC (1:1000, clone 30-F11, Biolegend, 103108), PI (1:100) and cell hashing antibodies (1:1000, #1 for heart, #2 for blood, Biolegend) and incubated for 30min protected from light. For TAC samples, PBMCs were purified as above, cardiac non-myocytes were directly used for cell sorting. Both cell fractions were stained with cell hashing antibodies (1:1000, #2 for heart, #3 for blood, Biolegend) and Propidium iodide (1:100, Thermo) and incubated for 30min. After washing, live cells of all samples were sorted on a Sony cell sorter SH800S, heart and blood cells of each genotype and condition were pooled and run on a Chromium controller using the Chromium Next GEM Single Cell 3ʹ v3.1 Library & Gel Bead Kit (10X Genomics) with a target of 10,000 cells per pool. Libraries were cloned according to the 10x Genomics user guide with feature barcode technology for cell surface protein (RevD) and Biolegend’s protocol for amplification of the cell hashing libraries. Libraries were pooled according to the recommendations of 10x genomics protocol and sequenced on a NovaSeq6000 (Illumina).

### Purification of primary cells and cardiac macrophages by F4/80 MACS and cell sorting for RNA-seq

For a total myocardial cell sample, 100µl of the raw cell suspension of primary cells from adult wildtype mouse hearts were washed twice with PBS and lysed in Trizol (ThermoFisher) for subsequent RNA preparation. For the purification of cardiac macrophages, the cell suspension was filtered through cell sieves of 70µm and 40µm of size and centrifuged at 100 *x g* for 3min at 4°C to completely remove cardiac myocytes. All following steps were performed on ice/at 4°C. The supernatant (non-myocytes) was washed twice in magnetic-activated cell sorting (MACS) buffer (PBS + 0.5% biotin-free BSA + 2mM EDTA). MACS was performed in a two-step process using anti-F4/80-biotin coupled antibody (clone BM8, Biolegend, 123106) and Streptavidin microbeads (Miltenyi Biotec) on LS columns (Miltenyi Biotec). The eluted macrophage fraction was resuspended in a final volume of 200µl MACS buffer and incubated 10 min with Fc block (1:50, BD Pharmingen). 200µl of staining solution containing flow cytometry antibodies (anti-CD45-PECy7 (1:100, clone 30-F11, Invitrogen, 25-0451-82), anti-CD64-PE/Dazzle594 (1:50, clone X54-5/7.1, Biolegend, 139320), anti-CD11b-PE (1:100, clone M1/70, BD Pharmingen, 557397)) was added and incubated for 30min in the dark. Cells were washed twice with MACS buffer and sorted on a Biorad S3 cell sorter. CD45+/CD64+/CD11+ macrophages were collected (between 100k-170K cells), washed with PBS and lysed in Trizol (Invitrogen) for RNA preparation.

### Fibrosis staining

Paraffin sections (8 μm) of ventricular myocardium were stained with Sirius red and Fast green. Images of the whole heart were taken with a 10x objective using a AxioObserver.Z1 (Zeiss) motorized scanning-stage microscopy (130 x 85; Märzhäuser). Fibrosis of the left ventricular myocardium was quantified as the ratio of signals from Sirius red to Fast green in each section using the Metamorph software (Molecular Devices).

### CD68 staining

Cryosections (5 μm) of ventricular myocardium were fixed with methanol for 10 min at −20°C, washed thoroughly with tap water and treated with acetone for 1 min at −20°C. Sections were air dried and subsequently blocked with 5% goat serum in PBS for 1 h. Sections were stained with anti-mouse CD68 (1:100, clone FA-11, AbD Serotec, MCA 1957) at 4°C overnight. The sections were washed three times with PBS + 0.05 % Tween 20 and stained with anti-rat AF564 (1:100, Invitrogen, A11007) and SYTOX™ Green (1:1000, Life technologies, S7020). Images were acquired by confocal microscopy using 63x glycerol objective (Leica SP5) and analyzed using Metamorph (Molecular Devices).

### Generation of knockout macrophages

Conditionally immortalized hematopoietic progenitor cells were generated from a R26-Cas9-eGFP transgenic mouse line^41^ as previously described^40^. For inducing CRISPR-Cas9 mediated knockout in Hoxb8-FL cells, cells were electroporated with an sgRNA pair targeting the *Schlafenlnc* locus (IDT: Alt-R® CRISPR-Cas9 sgRNA, 5’ cut site 5’-GAGGTTCCATCAGTCTGACA-3’, 3’ cut site 5’-GGCTTGGAAGTAGAAAACCG-3’), as control, a single sgRNA was used (5’-ACCCCTTCAGCTTCACCTCC-3’). Transfected cells were single-cell cloned by limiting dilution assay into 96-well plates and screened for knockout clones. CRISPR-directed deletions on *Schlafenlnc* were identified by a PCR-based strategy to amplify the genomic DNA of the clones. Single-cell clones carrying the induced deletions were expanded and frozen for further use. To do this, extraction of genomic DNA was performed with QuickExtract™ DNA Extraction Solution (Lucigen) according to the manufacturer’s instruction. Briefly, approximately 100,000 cells were lysed in 50µl of QuickExtract by incubation as follows: 65°C – 15min, 68°C – 15min, 98°C – 10min. 50ng of genomic DNA was then subjected to PCR amplification with two primer pairs to distinguish successful *Schlafenlnc* knockout clones from wildtype or heterozygous knockout. *Schlafenlnc*_KO_forward 5’-CAGCCCTGAGTCAGAACATAC-3‘ and *Schlafenlnc*_KO_reverse 5‘-TCCTTAGGGTGCACATCTACT-3‘ were used for detection of a successfully cut site with a resulting amplicon length of 480bp, *Schlafenlnc*_WT_forward 5’-TCTGTCTCTACTCCCTCAACTC-3‘ and *Schlafenlnc*_WT_reverse 5‘-AAGGCATGTTCTGCTCCTAC-3‘ were used for detection of the wildtype allele/uncut gene locus with a resulting amplicon length of 320bp.

### Hoxb8-FL *Schlafenlnc* KO cell culture and differentiation

Hoxb8 progenitor cells were cultured in RPMI 1640 supplemented with 10% FCS, 1% Pen/Strep, 50 ng/mL FLT3L, 50nM b-mercaptoethanol and 1nM b-estradiol. FLT3L supernatant was obtained from a FLT3L-producing B16 melanoma cell line.

For differentiation of HoxB8 cells into macrophages, Hoxb8 progenitor cells were cultured in differentiation medium consisting of the culturing medium described above without b-estradiol and supplemented with 20% MCSF supernatant, which was obtained from a MCSF-producing L929 cell line. Differentiation was accomplished after 6 days.

### Generation of bone marrow-derived macrophages

Femurs and tibias from 6-12 week old C57BL/6N mice were prepared and bone marrow was isolated by flushing the bones with R10 medium (RPMI medium, 10% FCS, 1% antibiotics) with a syringe. To generate bone marrow derived macrophages, 3-4 mio. isolated bone marrow cells were seeded in a 10cm petri dish containing R10 medium supplemented with 20% MCSF supernatant. Medium was added on day 3 and exchanged on day 6 after seeding, macrophages were fully differentiated on day 8 and used for experiments.

### Cytoplasmic/nuclear fractionation

Cytoplasmic/nuclear fractionation was performed as described before^67^. Briefly, bone marrow-derived macrophages (BMDMs) were washed two times with cold PBS and harvested. 10% of the cells was transferred into a new tube and used as a sample for the total gene expression. Cells were centrifuged at 200 *x g* for 10 minutes and the total expression sample was directly used for RNA isolation. The fractionation sample was resuspended in 200µl lysis buffer A (10 mM Tris pH=8, 140 mM NaCl, 1.5 mM MgCl2, 0.5% Nonidet P-40) and incubated for 5 minutes on ice. The lysate was centrifuged at 1000 *x g* for 3 minutes at 4°C. The supernatant containing the cytoplasmic fraction was directly used for RNA isolation through lysis with Trizol. The sediment containing the nuclear fraction was washed two times with lysis buffer A and one time with lysis buffer B (Buffer A supplemented with 1% Tween-40, 0.5% deoxycholic acid) and afterwards resuspended in Trizol for RNA isolation.

### RNA cloning and sequencing

Libraries from primary cardiac macrophages and total myocardium as well as *Schlafenlnc*-deficient and control cell lines were generated using the TruSeq Stranded RNA Kit (Illumina) according to the manufacturer’s protocol, except for one step. Instead of using the SuperScriptII (ThermoFisher) as the reverse transcriptase, SuperScriptIII (ThermoFisher) was used at a reaction temperature of 50°C. Sequencing was performed on a HiSeq4000 platform (Illumina).

Libraries from fractionation samples were generated using the Stranded mRNA Prep, Ligation Kit (Illumina) according to the manufacturer’s instructions. Sequencing was performed on a NovaSeq6000 machine (Illumina).

### RNA *in situ* hybridization (RNAscope)

Bone marrow derived macrophages were seeded on a removable 8-well chambers slide (Ibidi) and RNA *in situ* hybridization was performed using the RNAscope Fluorescent Multiplex Reagent Kit (Advanced Cell Diagnostics (ACD)) according to manufacturer’s instructions. A custom designed RNA probe by ACD was used for detection of *Schlafenlnc* (ENSMUST00000143673.2). Following the RNA *in situ* hybridization, the slides were incubated with Phalloidin-AF647 (Abcam) for 60 minutes at room temperature. Subsequently, slides were washed three times with PBS and mounted using ProLong™ Gold Antifade Mountant (Invitrogen). Images were acquired using the confocal microscope Leica TCS SP II and the Leica Application Suite.

### Galectin-3 ELISA

Cell culture supernatant of differentiated Hoxb8 macrophages was collected and the amount of soluble Galectin-3 was quantitatively assessed using the Galectin-3 Mouse SimpleStep enzyme-linked immuno-sorbent assay (ELISA) kit (Abcam) according to the manufacturer’s instructions. The cell culture supernatant was diluted 1:10 in sample diluent before use.

### Chemotaxis

An IncuCyte® ClearView 96-Well Chemotaxis Plate (Sartorius) was coated on both sides of the membrane using 50 µg/mL Matrigel (Corning) at 37°C for 30 minutes. After coating, plates were incubated at room temperature for 30 minutes and afterwards washed with D-PBS. Differentiated HoxB8 macrophages were harvested and 2,000 cells were seeded in each well in differentiation medium containing 0.5% FCS. Cells were allowed to settle at the bottom of the upper well for 30 minutes. Subsequently, 100nM recombinant murine C5a (Pepro Tech) was added to the lower well of the ClearView migration plate. Cells were imaged every 2 h for 72 hours using an Incucyte ZOOM instrument and a 10x objective with chemotaxis plate settings. Chemotaxis was analyzed using the IncuCyte ZOOM software.

### Adhesion Assay

Ibidi black 96 well µ-Plate was coated with 20µg/mL Fibronectin (Sigma Aldrich) for 1 h at 37°C. 10,000 differentiated Hoxb8 macrophages were seeded in each well in differentiation medium and were incubated for 2 hours at 37°C in a 5% CO2 atmosphere. After the incubation, cells were stained using NucBlue Live ReadyProbes reagent (Invitrogen) and following the manufacturers’ instructions. Images of the Nuclei were taken before and after removing the non-adherent cells by washing three times with pre-warmed medium using an EVOS FL Auto 2 cell imaging system (Invitrogen). Cell nuclei were counted automatically using the MetaMorph analysis software (Molecular Devices).

### Northern blotting

Northern blotting was performed as reported previously^68^. Briefly, 10µg total RNA from fibroblasts (NIH/3T3) and *in vitro* differentiated macrophages were run on a 1.2% formaldehyde agarose gel, transferred to a Hybond-N+ membrane (Ge Healthcare) and UV-crosslinked. Probes were end-labelled using [gamma-^32^P]-ATP and purified using an Illustra ProbeQuant G-50 micro column (GE Healthcare). A mixture of two different probes was used per target. Probes were incubated over night with a pre-hybridized membrane, signal was accumulated using a phophoimager screen and detected using a Cyclone Plus (PerkinElmer). mGAPDH probe1 5’-CATGGACTGTGGTCATGAGCCCTTCCACAATGCCAAAGTTG-3’, mGAPDH probe2 5’-GAATTTGCCGTGAGTGGAGTCATACTGGAACATGTAGACCATGTAGTTGAGG-3’, Slfnlnc probe1 5’-CAGCTCTGCGTCTCTGCTCCTGTTCAGGTCCATCTTTCATCTCCG-3’, Slfnlnc probe2 5’-CAGTCGGGGTTCTGCCTCTTGATGTGGTTGTGCTAACCGGTGGGAC-3’.

### *In vitro* transcription of 3’-biotinylated *Schlafenlnc* sense or antisense RNA probe

The *Schlafenlnc* sense and antisense (control) sequence was amplified from a vector containing the full *Schlafenlnc* (*AI662270-202*) sequence using Phusion DNA polymerase and the primers stated below. RNA probes for the RNA pulldown were *in vitro* transcribed from the amplified sequence using T7 RNA polymerase for 2h at 37°C. The probes were purified using illustra™ MicroSpin™ G-50 columns (GE Healthcare) and afterwards biotin labelled using the Pierce™ RNA 3’ End Biotinylation Kit (Thermo Scientific) and following the manufacturers’ instructions. RNA was extracted using Trizol and the quality and purity of the RNA probes was analyzed by Sanger Sequencing and Tapestation (Agilent).

T7_Slfnlnc_sense_fwd 5’-TAATACGACTCACTATAGGGAAAGTCGCCTGGGGCCTC −3’, T7_Slfnlnc_sense_rev 5’-AATGAGCAAGATCCAACAACTTT −3’, T7_Slfnlnc_antisense_fwd 5’-TAATACGACTCACTATAGGGAATGAGCAAGATCCAACAACTTT −3’, T7_Slfnlnc_antisense_rev 5’-AAAGTCGCCTGGGGCCTC −3’.

RNA probes were folded directly before use by incubating 150ng of biotinylated purified RNA in 75µl folding buffer (20mM Tris/HCl (pH 7.5), 100mM KCl, 10mM MgCl_2_ and Rnase inhibitor (New England Biolabs)) at 90°C for 2 minutes followed by incubation on ice for 3 minutes and finally incubation at room temperature for 25 minutes.

### RNA pulldown and mass spectrometry

10 million differentiated bone marrow-derived macrophages (isolated from C57BL/6N mice) were lysed in 500µl lysis buffer (50mM Tris HCl (pH8), 150mM NaCl, 0.5% (v/v) NP-40 with Protease (cOmplete, Roche) and Rnase inhibitor) for 30 minutes on ice. The lysate was centrifuged for 30 minutes at 17.000x*g* and 4°C. After the centrifugation, the supernatant was transferred into a new tube and 400µl dilution buffer (20mM Tris/HCl (pH 7.4), 150mM NaCl, 2mM EDTA (pH 8.00), 0.5% Triton X-100 with Protease and Rnase inhibitor) was added. 20µl streptavidin beads (Dynabeads M-270 Strepavidin, Invitrogen) were washed and resuspended in 100µl dilution buffer. The streptavidin beads were added to the diluted lysate and incubated for 30 minutes at 4°C with rotation to preclear the supernatant. After the incubation time, the beads were removed from the mixture and an input sample was taken. 150 ng *in vitro* transcribed 3’-biotinylated *Schlafenlnc* sense or antisense (control) RNA probe was added to the sample and incubated overnight at 4°C with rotation. 80 µl streptavidin beads were washed three times with fresh dilution buffer and resuspended in 100µl dilution buffer. The beads were added to the sample and incubated for three hours at 4°C with rotation. After the incubation time, the sample was washed three times with 500 µl dilution buffer for 10 min at 4°C with rotation. The sample was washed one more time with 500 µl dilution buffer containing 200 mM NaCl for 10 min at 4°C with rotation. The sample was washed two more times with MS-washing buffer (10mM Tris pH 7.4, 150mM NaCl) and analyzed with mass spectrometry. Liquid chromatography / mass spectrometry (LC/MS) was performed on Thermo Scientific™ Q Exactive Plus equipped with an ultra-high performance liquid chromatography unit (Thermo Scientific Dionex Ultimate 3000) and a Nanospray Flex Ion-Source (Thermo Scientific).

For data analysis MaxQuant 1.6.17.0^76^, Perseus 1.6.1.3^77^ and Excel (Microsoft Office 2016) were used. N-terminal acetylation (+42.01) and oxidation of methionine (+15.99) were selected as variable modifications and carbamidomethylation (+57.02) on cysteines as a fixed modification. Intensity based absolute quantification values (iBAQ) were recorded. The mouse reference proteome set (Uniprot, April 2021, 55470 entries) was used to identify peptides and proteins with a false discovery rate (FDR) less than 1%. Reverse identifications and common contaminants were removed. Proteins were filtered to be identified at least 4 times (n=6) in one experimental group. Missing values were replaced by background values from normal distribution. Significant interacting proteins were determined by student’s t-Test.

### PURA Immunoprecipitation

50µl Dynabeads Protein G beads (Invitrogen) were washed with PBS and incubated with 1ml anti-PURA antibody supernatant (monoclonal rat IgG2-2a anti-PURA^48^, clone: 12D11) or control antibody (rat IgG2a, clone: eBR2a, Invitrogen) overnight at 4°C with rotation. After the incubation, the beads were washed two times with PBS and transferred to a fresh Eppendorf tube. Differentiated BMDMs (7-8 mio. cells per sample, isolated from C57BL/6N mice) were washed and resuspended in 2 ml lysis buffer (25mM Tris HCl (pH 7.4), 150mM KCl, 0.5% NP-40, 1mM EDTA (pH 8.0), protease (cOmplete, Roche) and Rnase inhibitor (NEB)). Cells were sonicated for 30 seconds (Bandelin electronic sonicator UW 3100, duty cycle 0.5, amplitude 30%, MS72 tip) and incubated for 30 minutes on ice. The lysate was centrifuged for 30 minutes at 17.000xg at 4°C and the supernatant was transferred to a fresh Eppendorf tube. An input sample was taken at this point. The lysate was added to the magnetic beads and incubated for 4 hours at 4°C with rotation. After the incubation, the beads were washed three times with washing buffer (50mM Tris/HCl (pH 7.4), 150mM KCl, 0.1% NP-40, protease and Rnase inhibitor). The PURA pull-down was confirmed with SDS-PAGE and Western blot and the *Schlafenlnc* RNA as well as a positive (*Tpt1*) and negative control (*Rpl32*) were measured by qPCR analysis.

### Quantitative real-time PCR

1 μg of total RNA was reverse transcribed with Protoscript II reverse transcriptase (NEB), both according to the manufacturer’s instructions using random hexamer primers. Quantitative real-time PCR (qPCR) was carried out with primers listed below, using the FastStart universal SYBR Green master mix (Roche). The sample volume of 12.5 μl contained 1x SYBR Green master mix, 5 pmol of each primer and 10 ng of cDNA. PCR was performed using StepOnePlus Real-Time PCR systems (Applied Biosystems).

Slfnlnc_qPCR fwd 5’-ATGACTGGTCCTGGTTTTCG −3’, Slfnlnc_qPCR fwd 5’-TCTGCCTCTTGATGTGGTTG −3’, mRPL32 fwd (housekeeping gene) 5’-ACATCGGTTATGGGAGCAAC −3’, mRPL32 rev (housekeeping gene) 5’-GGGATTGGTGACTCTGATGG −3’, Tpt1 fwd (positive control for PURA IP) 5’-AGTCACCGGTGTTGACATTG -3’, Tpt1 rev (positive control for PURA IP) 5’-GCTCTGCAGCTCCAGTCATA -3’.

### Data analysis

#### Single cell sequencing data analysis

Raw reads generated by single cell RNA sequencing data from sham^16^ and 6d after TAC (this study) was aligned against the mouse genome (mm10, Gencode annotation vM22^18^) using CellRanger 5.0.0 (10x Genomics). Cell that were called by CellRanger were subsequently analyzed using the R package Seurat v3.2^69^. After quality control (retaining cells with >500 or <3,600 detected genes, <10% mitochondrial cell content), 7,913 cells of the sham condition and 13,803 of the TAC condition were used for further downstream analysis. SCTransform^70^ was used to normalize and scale the data as well as remove unwanted variation (regressing out mitochondrial gene content and cell cycle score). SCTransform was then applied to identify variable feature which were used to integrate both data sets (“IntegrateData” function). By using the ElbowPlot function on Principal Component Analysis results, the number of relevant components was determined (set to 40) and used for embedding and visualization by Uniform Manifold Approximation and Projection (UMAP). Seurat’s “FindAllMarkers” function was used for determination of cell population marker genes.

For the comparison of clusterings with adjusted Rand index score, gene sets were defined as follows: all genes = all genes listed in the vM22 annotation of Gencode; protein coding genes = all genes with gene biotype “protein coding” in vM22; lncRNAs = all genes in vM22 lncRNA annotation of Gencode (gene biotype “lncRNA”+“TEC”); “expressed” gene sets = same definitions as above with filter for average expression >0 in at least one cell cluster; all variable genes = gene list generated by Seurat’s “SelectIntegrationFeatures”, default is 3,000 genes; subset of variable genes based on gene lists defined above. Next, clusterings with resolutions from 0.01 to 1.00 with 0.01 increments were iteratively calculated (“FindClusters” function) and cluster ID per cell was saved in a data frame. The adjusted Rand Index is a measure for the degree of similarity between two clusterings, for example. The R package mclust^71^ with inbuilt function “adjustedRandIndex” was used to compare all gene sets to the respective complete gene set (e.g. the subset of expressed lncRNAs to all expressed genes).

For calculating de-regulated genes after TAC, first Adaptively thresholded Low-Rank Approximation (ALRA)^72^ was performed in order to impute drop out gene expression values followed by using Seurat’s “FindMarkers” function to compare each cell population before and after TAC. For the analysis of scRNA-seq data of *Schlafenlnc*-deficient and wildtype animals, raw reads from basal and 6d after TAC from WT and KO were aligned against the mouse genome (mm10, Gencode annotation vM25^18^) using CellRanger 6.0.1 (10x Genomics) and analyzed using the R package Seurat v3.2^69^. Quality control was applied to basal (>200 or <5,000 detected genes, <10% mitochondrial cell content) and TAC (>200 or <4,000 detected genes, <10% mitochondrial cell content) samples. SCTransform and FindAllMarkers was applied as above to normalize, scale, integrate data and identify marker genes for cell populations. The integrated object was subset on all cell clusters expressing *Cd45* (Leukocytes). Deregulated genes were calculated using Seurat’s “FindMarkers” function to compare each cell population between *Schlafenlnc*-deficient and wildtype cells. GO Term analysis of the deregulated genes was performed using “DAVID 2021”^73^.

#### Cell-cell communication analysis

A subset of cells of interest from blood and heart of *Schlafenlnc*-decient and wildtype *Cd45*+ (*Lyve1* macrophages, *MHCII*-high macrophages, *Isg* macrophages, intermediate macrophages, classical monocytes and non-classical monocytes) was used for analysis of secreted signaling pathways within the CellChat database. The *Schlafenlnc*-deficient and wildtype objects were analyzed separately using CellChat^74^ and merged with “mergeCellChat”. The differential number of interactions between the cell clusters in *Schlafenlnc*-deficient vs. wildtype was analyzed and displayed using the CellChat function “netVisual_diffInteraction” and the differential signaling pathways were assessed using “rankNet”.

#### Estimating RNA velocity

To calculate the RNA velocity of single cells, the abundance of the spliced and unspliced RNA was quantified by running Velocyto CLI (v0.17.5) on CellRanger output BAM files and the corresponding gene annotation (GENCODE vM22). The resulting loom files were incorporated into the Seurat objects using SeuratWrapper R package (v0.3.0), and after integration and clustering, the objects were converted to AnnData format (h5ad) to be compatible for downstream analysis with the Python package scVelo (v0.2.5). Using scVelo, cell-to-cell transition probabilities were estimated and RNA-velocity was projected as streamlines on the UMAP embedding.

#### Identification of novel lncRNAs

Most steps of this pipeline were performed using the Galaxy platform^75^. Reads were mapped using RNA STAR aligner (Galaxy version 2.6.0b-2)^76^ using a built-in index based on Gencode vM22 annotation and otherwise default parameters. StringTie (Galaxy Version 1.3.3.2)^77^ was used to predict novel transcripts (strand information RF, reference file Gencode vM22, average read length 90). StringTie merge (Galaxy version 1.3.6) with default parameters was used to merge all new annotations together, including Gencode annotation vM22 as reference annotation. Novel genes were extracted by Bedtools Intersect tool (Galaxy version v2.29.0)^78^, intersecting the merged annotation with the Gencode vM22 annotation. Transcripts were extracted with the “Extract features” tool (Galaxy 1.0.0) and analyzed for coding potential with FEELnc (Galaxy version 0.1.1.1)^30^. FEELnc produces a gene transfer format (gtf) annotation file containing only transcripts classified as long non-coding RNAs. “cat” command was used to merge this lncRNA annotation to Gencode vM22 reference annotation file to create reference annotation file with additional novel lncRNA genes. All RNA-Seq libraries were measured again against this new transcriptome using StringTie’s “count” option. For visualizing coding probability, the python version of the Coding-Potential Assessment Tool (CPAT, v2.0.0)^79^ was run on the fasta sequences of the novel lncRNAs ($ cpat.py -g novel_lncRNAs_gffread.fasta -d Mouse_logitModel_CPAT.RData -x Mouse_Hexamer_CPAT.tsv -o output_novel_lncRNAs.txt). GraphPad Prism 8 was used to calculate the frequency distribution in percentages. For plotting transcript lengths, the length determined in the CPAT output was used and again transformed to frequency distribution in Prism. For the categorization of interactions between lncRNAs and other genes, the classifier output of FEELnc was used. Interactions with the same gene were discarded, for known lncRNAs only genes expressed predominantly in cardiac macrophages were considered (log_2_ fold change >1). To compute differential gene expression analysis, DESeq2 package was used (Galaxy version 2.11.40.2)^80^, outputting also a normalized counts file. Normalized counts served as input for clustering and heatmap visualization by ClustVis R package (v. 0.0.0.9)^81^.

#### RNA-Seq analysis of Hoxb8 Schlafenlnc-deficient macrophages

Fastq reads were mapped using RNA STAR aligner (Galaxy version 2.6.0b-2)^76^ using a built-in index based on Gencode vM22 annotation and otherwise default parameters. StringTie (Galaxy Version 1.3.3.2)^77^ count option was used to quantify gene expression using Gencode vM22 as reference annotation. DESeq2 was used to calculate differential gene expression (Galaxy version 2.11.40.2)^80^. The normalized counts file was used for heatmap visualization with ln(x) transformation of the top 20 down-regulated genes (ClustVis R package^81^).

#### Neighboring genes analysis and Gene Ontology Analysis

Up- or downregulated lncRNAs during pressure overload (p-value<0.05) in either macrophage, fibroblast or endothelial cell populations were analyzed for neighboring genes using FEELnc (Galaxy Version 0.2^30^) and filtered for antisense protein coding genes. Gene ontology analysis for these gene sets for each cell population was performed using the web-based tool “DAVID 2021”^73^. Neighboring genes of known and novel lncRNAs were also identified using FEELnc (Galaxy Version 0.2^30^).

Biological processes associated with the differentially regulated genes in *Schlafenlnc*-deficient macrophages (log2FC > 1 or < -1, FDR < 0.05) were determined using ClueGO application (v2.5.8)^82^ ran within Cytoscape (v3.8.2)^83^ network analysis tool (p-value cutoff = 0.01, Min GO level = 4, Max GO Level = 10, Number of Genes = 2, Min. Percentage = 4.0).

#### Kipoi RBP-eclip analysis (Protein interactors analysis)

Model training data: To access the binding probability of known RBPs on *Schlafenlnc*, a pretrained deep neural network (DNN)^34^ on eCLIP-seq data^35^ was downloaded from the Kipoi repository^84^. In order to test the binding probability of the 101 proteins on *Schlafenlnc*, a bed file containing the coordinates of *Schlafenlnc*’s 4 exons (ENSMUST00000143673.2 on Ensembl) was obtained. Then for each exon, we took sliding windows of 101nt length resulting on a new bed6 file where each row is of 101nt length. Kipoi builds on environments, making the sharing and loading of models easy. We started our analysis by generating an environment with all the needed dependencies. Afterwards for each protein of interest, we programmatically obtained the dataloader, load the fasta sequence for GRCm38.p6, load the GENCODE annotation for mouse version vM22^18^, and a bed file with the pre-extracted 101nt length sliding windows and ran the predict function with default arguments (considering all the genomic landmarks also for non-protein coding genes). Finally, for each protein of interest a csv file was generated, containing the following info for each 101nt sequence considered: the chromosome, the starting position, the end position, a sequence ID, a score as in the bed6 file, the strand and the binding score [0 (no binding probability), 1 (high binding probability)] of the protein of interest on that window.

### Software list

Leica Application Suite Advanced Fluorescence, version 2.7.9723

Invitrogen EVOS FL Auto 2 Software

CellRanger, version 5.0.0

Seurat, version 3.2

RNA STAR aligner, version 2.6.0b-2

StringTie and StringTie merge, version 1.3.3.2 and 1.3.6

Bedtools Intersect tool, version v2.29.0

“Extract features” tool, version 1.0.0

FEELnc, version 0.1.1.1 (Galaxy) and version 0.2

Coding-Potential Assessment Tool, CPAT, version 2.0.0

DESeq2, version 2.11.40.2

ClustVis R package, version 0.0.0.9

Cytoscape, version 3.8.2

ClueGO, version 2.5.8

MetaMorph Offline, version 7.10.1

Prism 8 for Mac OS, version 8.4.1

Adobe Illustrator for Mac OS, version 24.2.1

R studio, version 3.6.3

Microsoft Office for Mac

### Statistical Analysis

Statistical analysis was performed using GraphPad Prism 8. To test for outliers Grubbs test (alpha=0.05) was used. Data distribution and the difference between variances were tested using the Shapiro-Wilk normality test. For statistical analysis of two groups, unpaired t-test was performed for normal distributed samples, otherwise Mann-Whitney test was used. For more than two groups, two-way ANOVA was performed. For chemotaxis experiments, area under the curve (AUC) was calculated and statistical significance was tested using unpaired t-test. Significance is depicted as *p < 0.05, **p < 0.01, ***p < 0.001, ****p < 0.0001.

## ACKNOWLEDGEMENT

We thank Lucia Koblitz for expert support with primary cell preparations, Julia Kerler and Anton Bomhard for expert assistance in animal surgeries and Dr. Elisabeth Graf for assistance with RNA-seq. We thank Dr. Daniel Andergassen and Dr. Bernhard Laggerbauer for critical reading and helpful discussions.

## Author contributions

A.D. and S.E. designed the study and wrote the manuscript. A.D., L.A. D.E., S.B., E.T., K.H., N.P., A.U. and I.W. performed the experiments or analyzed the resulting data. M.B. provided the *Schlafenlnc* knockout mouse. R.P.B., J.G., P.K., M.S.L., L.M., T.M., D.R., H.B.S., M.S.-S., C.S., D.N. and T.W. provided experimental models, instruments and crucial expertise.

