## Supplemental Data Figures for "The lncRNA landscape of cardiac resident macrophages and identification of *Schlafenlnc* as a regulator of macrophage migratory function"

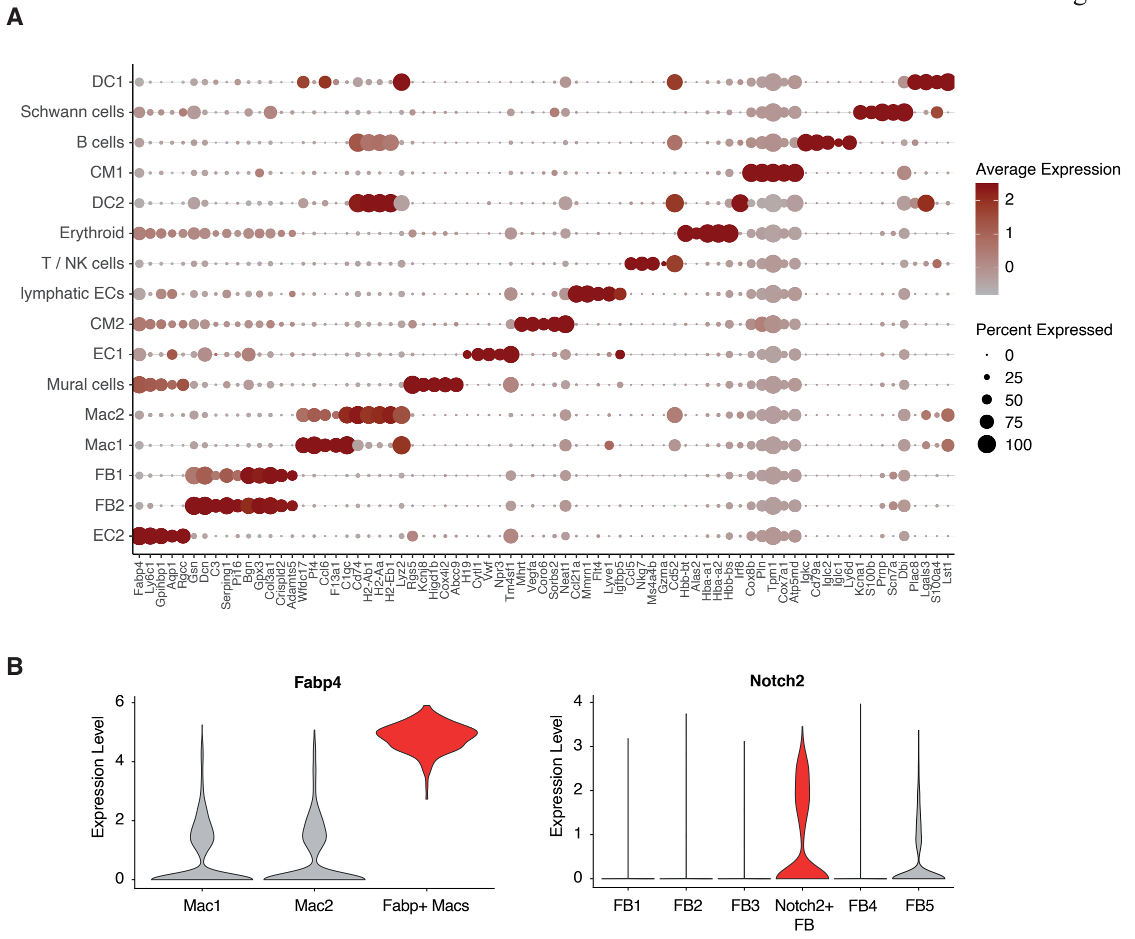


**Extended Data Figure 1:**

(A) Bubble plot for the top5 marker genes (all genes) in each cell population. Bubble size corresponds to percentage of cells the gene is expressed in, the color indicates the level of expression. (B) Expression of *Fabp4* and *Notch2* in all macrophage or fibroblast subclusters, respectively.

Extended Data Figure 2

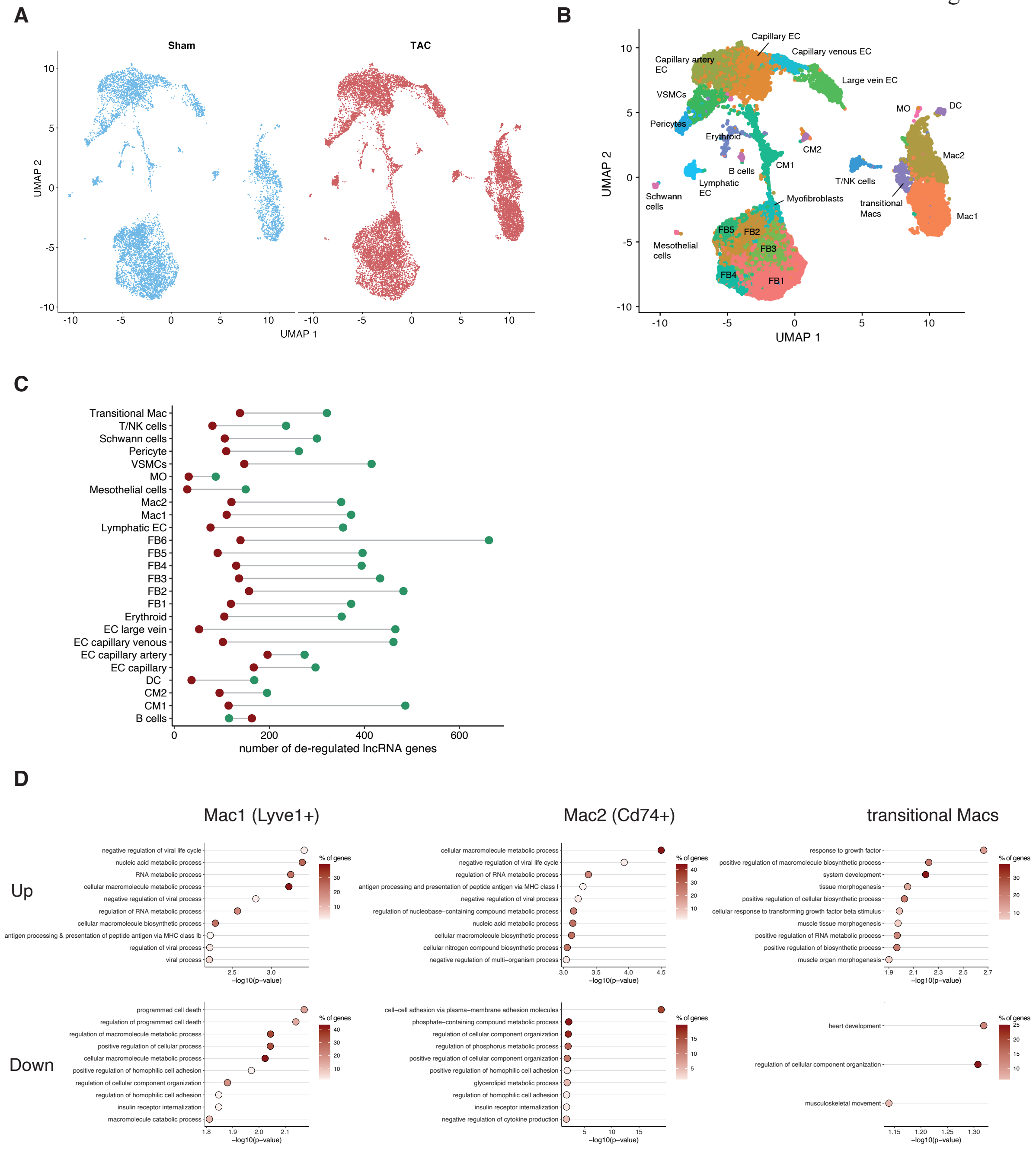


**Extended Data Figure 2:**

(A) Umap projection of murine myocardium after Sham or TAC as measured by scSeq. (B) Umap projection of murine myocardium with cell populations colored by type as indicated by the label. (C) Number of de-regulated lncRNAs after TAC (TAC vs. Sham) per cell cluster. Red dot signifies number of down-regulated genes, green dot shows number of up-regulated lncRNA (p-Val < 0.05). (D) Gene ontology term analysis of genes located antisense to down- (lnFC<0) or upregulated (lnFC>0) lncRNAs during TAC in different macrophage subpopulations.

Extended Data Figure 3

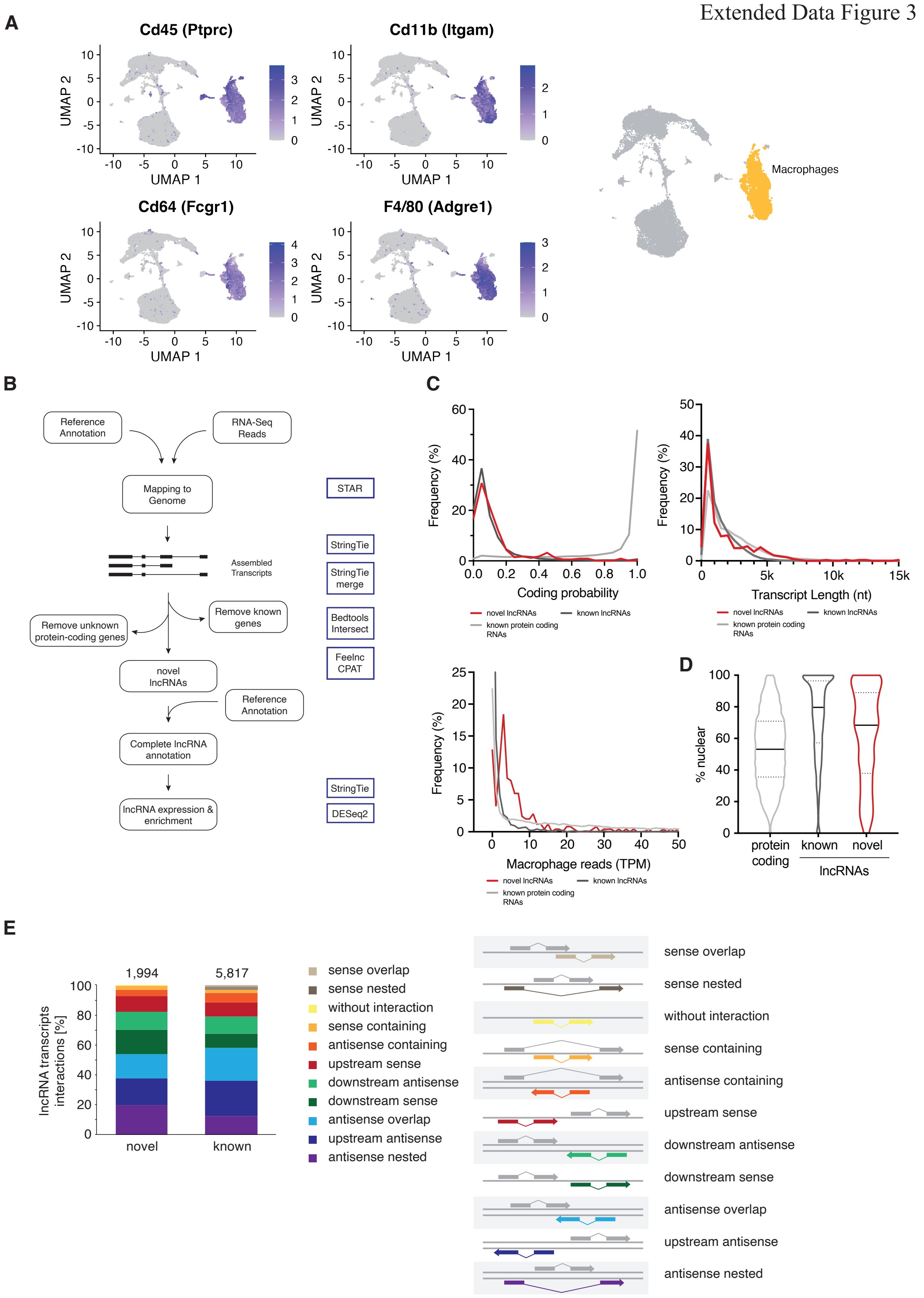


**Extended Data Figure 3:**

(A) FeaturePlots showing the expression of macrophage marker genes CD45, CD64, CD11b and F4/80. (B) Workflow for identification of novel lncRNAs. (C) Novel lncRNAs (shown in red) were analyzed for their coding potential using CPAT, their length and expression strength, comparing them to known lncRNAs (black) and known mRNAs (grey). All data presented as frequency distributions in percentages. (D) Distribution of the intracellular localization (cytoplasmic/nuclear) of mRNAs, known and novel lncRNAs as determined by fractionation of bone marrow-derived macrophages and subsequent RNA-Seq. Black line indicates median value, dotted lines quartiles. (E) Analysis of lncRNA interaction in a genomic context using FeeInc[21]. LncRNAs (separated into novel and known lncRNAs) were tested in which orientation they lie within 10kb of their start and end. Right panel depicts the categories graphically.

Extended Data Figure 4

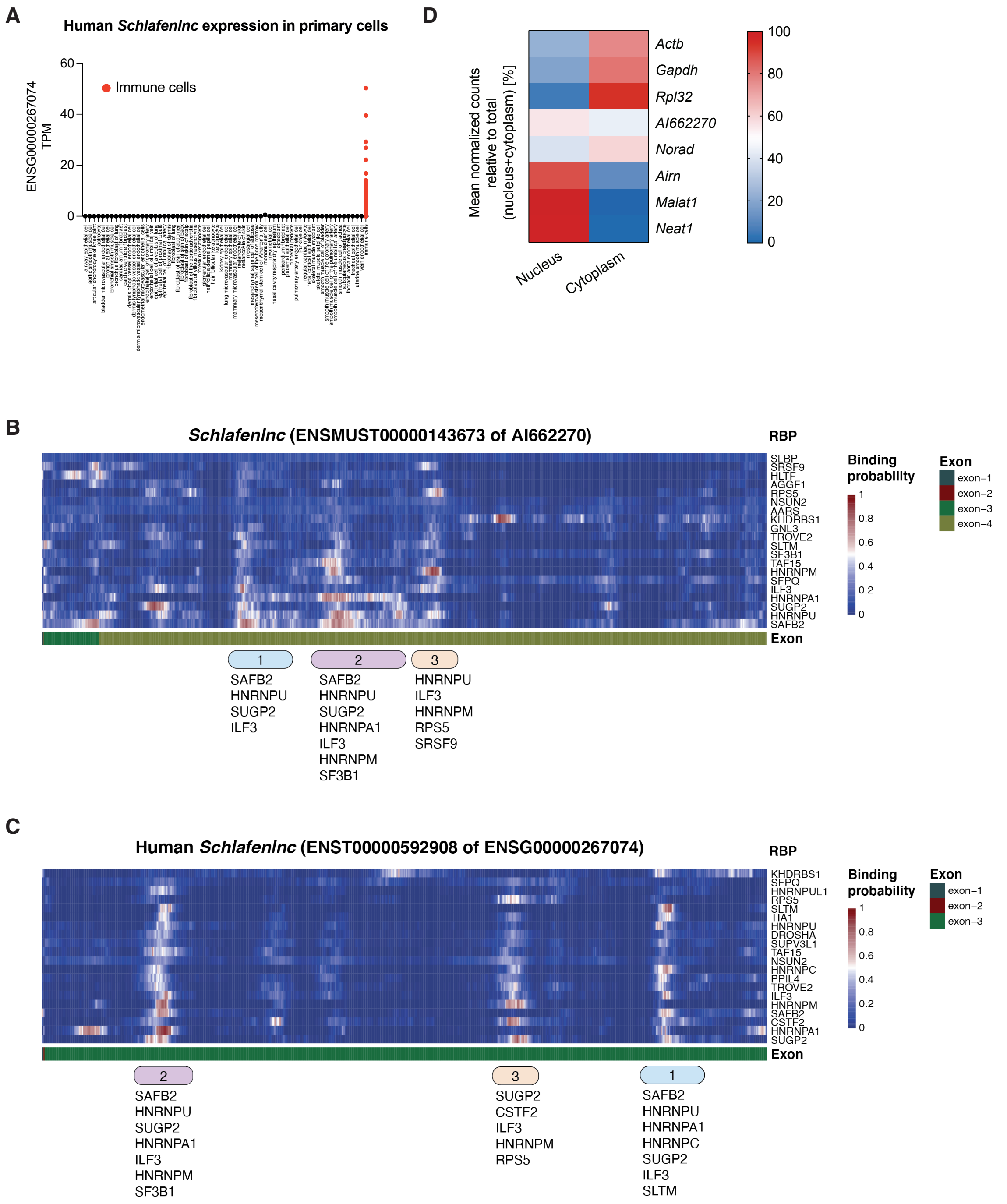


**Extended Data Figure 4:**

(A) Expression profile of human *Schlafenlnc* (*ENSG00000267074*) in transcript per million (TPM) in 205 data sets of primary cells in Encode database. Expression values were extracted using the RNA-Get functionality of the Encode website. (B) Distribution pattern of lncRNAs or mRNAs known to localize to the nucleus (*Malat1*, *Neat1*, *Airn*), cytoplasm (*Gapdh*, *Rpl32*, *Actb*) or both (*NORAD*) as validation for a successful fractionation. (C) Analysis of potential RNA binding protein (RBP) binding sites within *Schlafenlnc*. 101 RBPs were tested on the sequence of *Schlafenlnc* using the Kipoi RBP-eclip trained deep neural network. The sequence was binned into 101 nucleotide long windows and then tested for binding probability (ranging from 0 to 1). The top 20 strongest binders are shown here. Three clusters are highlighted and numbered as they represent potential binding hot spots. (D) Same analysis as in (C) using the sequence of human *Schlafenlnc* (*ENST00000592908*). Same cluster as in (C) are highlighted as well.

Extended Data Figure 5

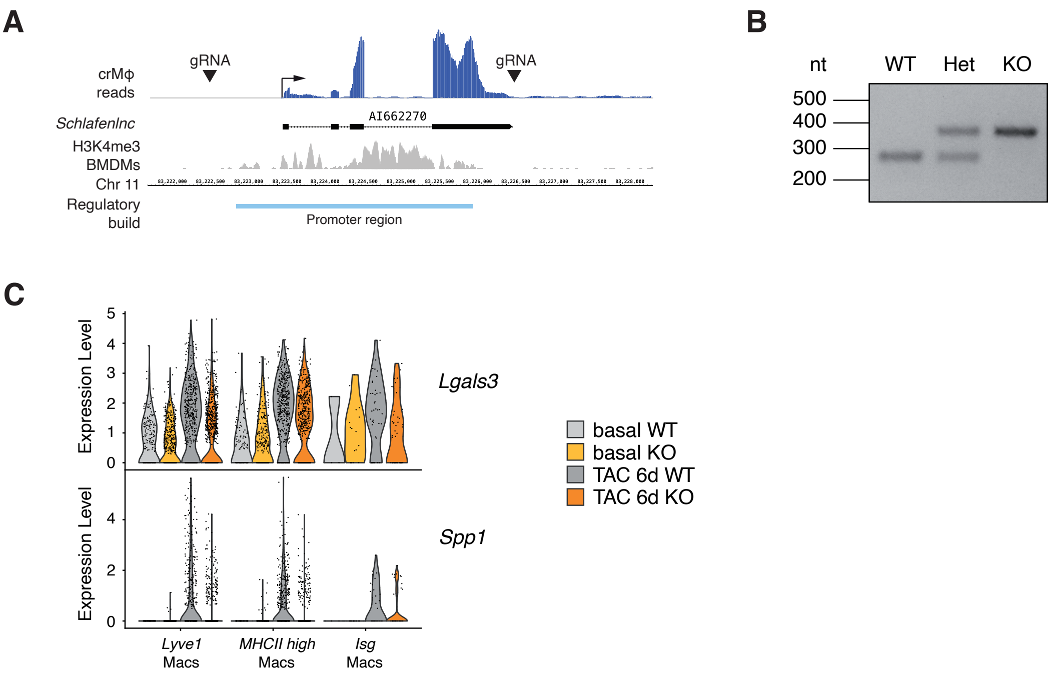


**Extended Data Figure 5:**

(A) Genomic locus of *Schlafenlnc* (*AI662270*) indicating induced cut sites to generate the knockout mouse line (black triangles). (B) Exemplary genotyping PCR, higher band (around 400bp) indicates successful knockout. (C) Expression levels of *Lgals3* (Galectin-3) and *Spp1* (Osteopontin) in macrophage population in WT and *Schlafenlnc*$^{-/-}$ cells.

Extended Data Figure 6

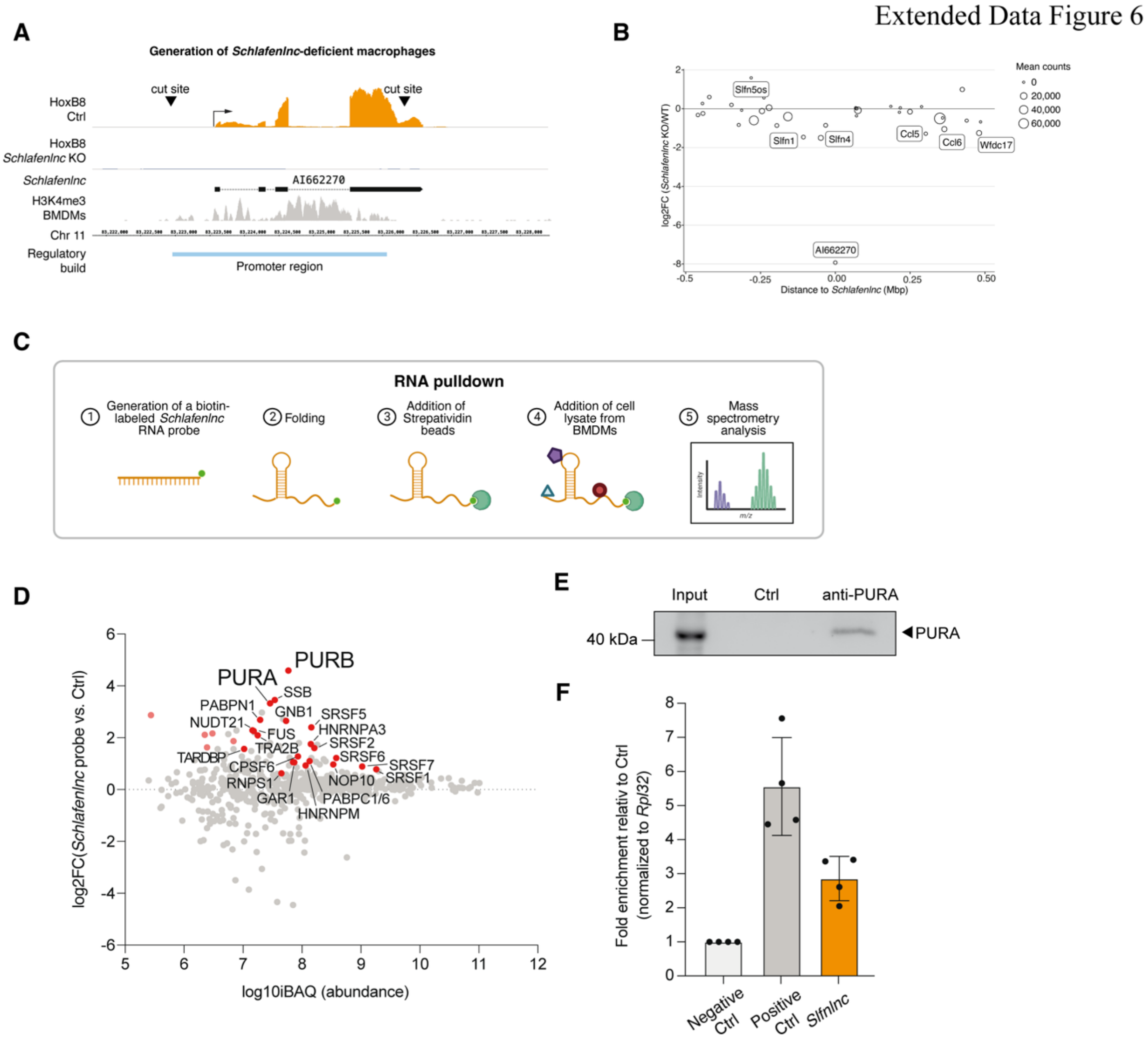


**Extended Data Figure 6:**

(A) Genomic locus of *Schlafenlnc* (*AI662270*) indicating induced cut sites to generate knockout Hoxb8-FL cells (black triangles). (B) Changes in gene expression in the vicinity of Schlafenlnc (+/- 0.5 Mbp). Bubble size indicates mean expression over all samples (WT and KO). Genes with |log2FC| > 1 are labelled by name. (C) Scheme of the workflow for the RNA pulldown experiment with an *in vitro*-transcribed, biotin-labelled *Schlafenlnc* probe. (D) RNA pulldown analysis, significantly enriched proteins on the *Schlafenlnc* probe are shown in red. (E) Western blot analysis of the input sample, the control sample, and the pulldown sample (anti-PURA) of the PURA pulldown experiment. (E) qPCR measurement of the PURA pulldown. Negative control = *Rpl32*, positive control = *Tpt1*. Data is normalized to the Ctrl sample and displayed as mean ± SEM.
